# The Pump-Leak/Donnan ion homeostasis strategies of skeletal muscle fibers and neurons

**DOI:** 10.1101/2020.11.20.391813

**Authors:** Béla Joos, Catherine E Morris

## Abstract

Skeletal muscle fibers (SMFs) and neurons are low and high duty-cycle excitable cells constituting exceptionally large and extraordinarily small fractions of vertebrate bodies. The immense ClC-1-based chloride-permeability (P_Cl_) of SMFs has thwarted understanding of their Pump-Leak/Donnan (P-L/D) ion homeostasis. After formally defining P-L/D set-points and feedbacks, we therefore devise a simple yet demonstrably realistic model for SMFs. Hyper-stimulated, it approximates rodent fibers’ ouabain-sensitive ATP-consumption. Size-matched neuron-model/SMF-model comparisons reveal steady-states occupying two ends of an energetics/resilience P-L/D continuum. Excitable neurons’ costly vulnerable process is Pump-Leak dominated. Electrically-reluctant SMFs’ robust low-cost process is Donnan dominated: collaboratively, Donnan effectors and [big P_Cl_] stabilize V_rest_, while SMFs’ exquisitely small P_Na_ minimizes ATP-consumption, thus maximizing resilience. “Classic” excitable cell homeostasis ([small P_Cl_][big I_Naleak_]), *de rigueur* for electrically-agile neurons, is untenable for vertebrates’ (including humans’) major tissue. Vertebrate bodies evolved thanks to syncytially-efficient SMFs using a Donnan dominated ([big P_Cl_][small I_Naleak_]) ion homeostatic strategy.

## INTRODUCTION

### Ion Homeostasis is a Pump-Leak/Donnan (P-L/D) feedback process

Ion homeostasis is the energy-consuming feedback process by which a cell maintains its steady-state volume (Vol_cell_), membrane potential (V_m_) and ion concentrations ([ion]_i_), returning autonomously to steady-state after ionic perturbations by sensing a change and transiently adjusting its fluxes and energy consumption till steady-state is restored. In the classic feedback control situation, thermostatic regulation of room temperature, a temperature sensor/effector system alters the output of a heat source in proportion to deviations from a chosen temperature set point. As per **Figure 1A** and legend, ion homeostatic feedback too, has sensor/effector elements and a set point. For any particular cell-type, the ion homeostasis set point is a steady-state collection --{Vol_cell_, V_m_, [ion]_i_}--“chosen” by evolution. Evolution furnishes that cell-type with: a particular quantity (Usher-Smith et al, 2009) of anionic Donnan effectors, an H_2_O-preamble membrane of a particular area (C_m_) equipped with particular densities of Na^+^, K^+^ and Cl^-^ permeant pathways (P_Na_, P_K_, P_Cl_), and electrogenic (3Na^+^/2K^+^-ATPase) pumps sufficient to sustain steady-state and restore it after perturbations of the size expected for that cell-type. Quantitative cell-type specific differences for these components account for different ion homeostatic set points. The minimal components for self-regulating Pump-Leak/Donnan (P-L/D) ion homeostasis are depicted in **Figure 1B**. In virtually all cells, ion homeostasis will be non-minimal since most deploy additional ion transporters and channels to address nuanced physiological requirements (Fraser and Huang 2004), but these components tune rather that cause P-L/D ion homeostasis and so are addressed here only peripherally.

**FIGURE 1.**
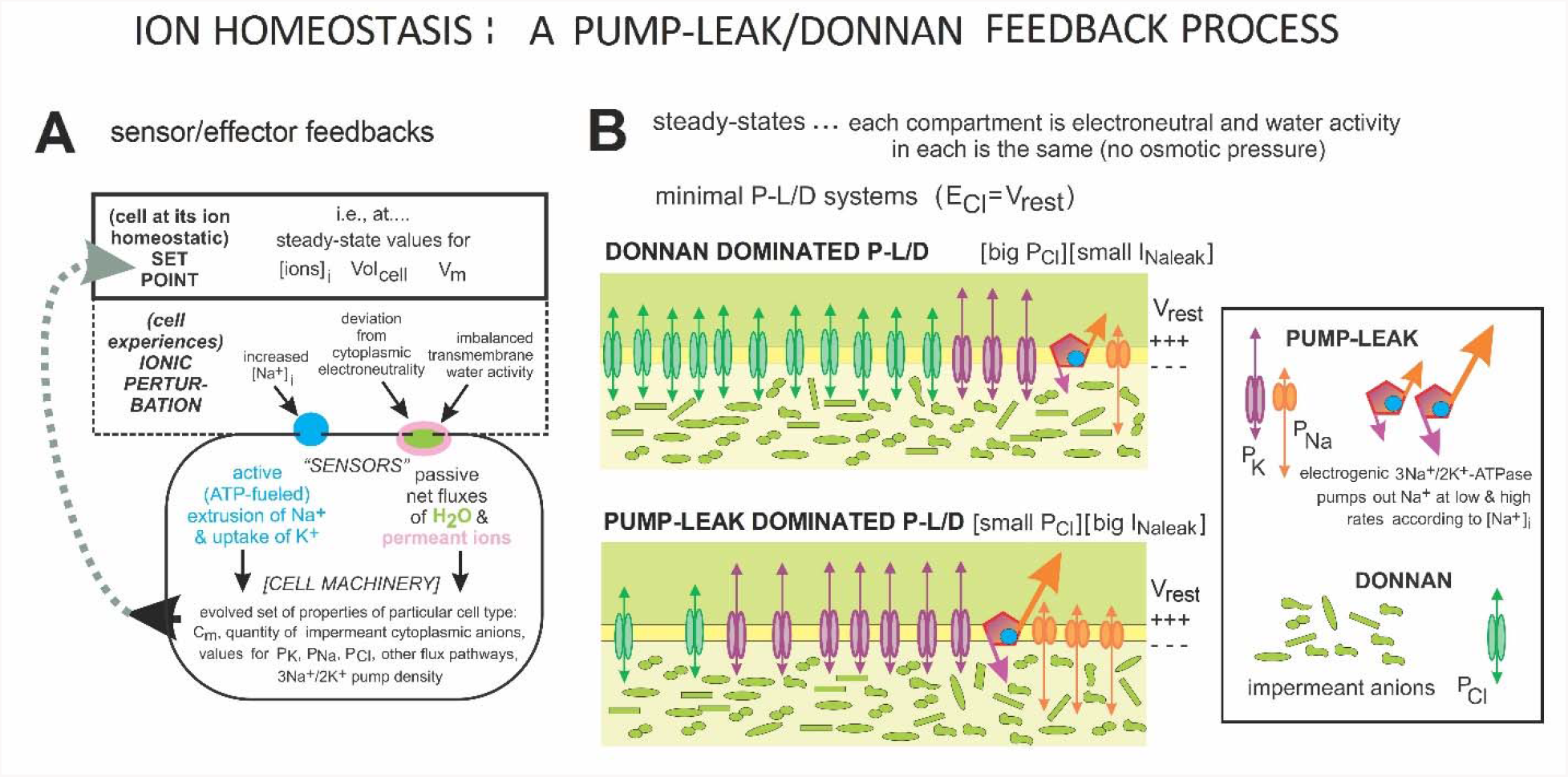
Fundamentals. **A**. **Ion homeostasis as an autonomous Pump-Leak/Donnan (P- L/D) feedback process with two sensor/effector mechanisms** P-L/D systems cannot sense deviations from their “set point” values. The otherwise blind-to-system-values 3Na^+^/2K^+^-pump functions as a sensor/effector by actively extruding Na^+^ faster (thus hyperpolarizing V_m_) when [Na^+^]_i_ is higher (as per **Figure 9Ai,ii**) while the (Donnan-effectors)/P_Cl_ combination, via thermodynamics/electrostatics, functions as a sensor/effector that prevents osmo-imbalance and maintains compartment neutrality. (The weak V-dependence of Na/K-pumps (Rakowski et al 1997) is not modeled here). **B**. **Minimal P-L/D systems at steady-state** Implied but not depicted: **1)** the membrane has an H_2_O-permeable bilayer of capacitance, C_m_, that can store charge until ∼V_m_±200 mV, **2)** compartments are neutral, osmo-balanced and iso-potential, **3)** the cellular surface-area-to-volume (SA:V) ratio handles normal swelling perturbations without bilayer-rupture, **4)** ATP is available ad libitum, **5)** average Donnan effector valence=-1, **6)** the exterior is infinite/invariant. V-gated channels are not depicted because normally they do not contribute to steady-state. Impermeant cytoplasmic anions (mostly proteins) act as Donnan effectors; Donnan effectors passively influence the transmembrane distributions of permeant ions and H_2_O. Animal cells’ electrogenic 3Na^+^out/2K^+^in-ATPases cycle faster at ↑[Na^+^]_i._ ([K^+^]_e_ is invariant here, so pump sensitivity to [K^+^]_e_ is not modeled). At ion homeostatic steady-state, the hyperpolarizing pump-flux precisely counterbalances in/out fluxes across P_Na_ and P_K_. For these minimal systems, V_m_=V_rest_=E_Cl_ because in/out anion flux across P_Cl_ balances at V_rest_. In minimal systems, moreover, the absolute quantity of Donnan effectors determines resting Vol_cell_ (Usher-smith et al 2009) (MN-CD vs CN-CD offers a simple minimal vs non-minimal comparison; see **Table 1** footnotes(**\*\***)). For self-regulating P-L/D ion homeostasis, these are the minimal components. In the top and bottom minimal systems, P_Na_:P_K_:P_Cl_ ratios differ, as does the steady-state pump-rate (implied by arrow-size). For the steady-state (net fluxes=0) at top, influx/efflux is predominantly anions (Cl^-^) passively balanced by Donnan effectors, and at bottom, influx/efflux is predominantly cations actively counterbalanced by [Na^+^]_i_-sensitive energy-consuming pump work. The top P-L/D ion homeostatic steady-state we describe as “Donnan dominated” ([big P_Cl_][small I_Naleak_]), the lower one as “Pump-Leak dominated” ([small P_Cl_][big I_Naleak_]).

### Evolutionary context for muscle and nerve ion homeostatic strategies

Skeletal muscle is the human body’s biggest tissue (∼40%; Janssen et al 2000). This tissue’s fundamental ion homeostatic mechanisms evolved to meet the needs of our early vertebrate ancestors. Their bodies comprised, perhaps, 40-60% skeletal muscle fiber (SMF) (see **Table 1**). Then, as now, SMFs would have been low duty-cycle excitable cells (i.e., at V_rest_ except during brief, infrequent action potentials (APs)). Also then as now, vertebrate neurons would have been high duty-cycle electrically-agile input-summing cells tasked by the whole-organism with overseeing sensory processing and controlling the timing of muscle contractions. For those ancestral vertebrates, a low-cost SMF resting-state would have been an enormous boon, while, given the tiny body portion devoted to brain in early vertebrates (**Table 1**), heavy ion homeostatic spending on neurons would have been even more tolerable than (say) in modern mammals. That ancient tissue-mass difference plus the starkly different electrophysiological lifestyles of skeletal muscle and neurons likely governed the evolution of the distinctive ion homeostatic strategies we identify here as operative in modern mammals (and modern amphibians, etc).

**Table 1.** Pump-Leak/Donnan models for excitable cells

|  | *Model name:*<br>*Item:* | **SM-CD** | **WD-CD** | **CN-CD** |
| --- | --- | --- | --- | --- |
| 1 | ***P-L/D model***<br>*an “ion homeostatic unit”*<br>*(i.e., same membrane $C_m$*<br>*for each) depicting:---* | one “slice” of the many needed<br>to comprise a multinucleate<br>skeletal muscle fiber<br>(explained in **Figure 8C,D**) | a <u>counterfactual</u><br>computational “tool”<br>(a Pump-Leak dominated<br>analog of SM-CD) | a roundish soma of a<br>cortical neuron in brain-slice<br>(Dijkstra et al 2016)<br>(termed CN-CD here) |
| 2 | ***in ancestral vertebrates*** | SMFs >40% of body mass* | --- | brain <0.2% of body mass* |
| 3 | ***in humans*** | SMFs ~40% of body mass<br>(e.g. Rolfe et al 1997) | --- | brain ~2% of body mass<br>(Mink et al 1981) |
| 4 | ***“electrophysiological***<br>***life-style” served*** | low excitability<br>low duty-cycle,<br>“off-on-off” AP signal | (same as SM-CD) | high excitability,<br>high duty-cycle,<br>complex nuanced $V_m$ signalling |
| 5 | ***dominant st-state fluxes*** | ANIONS ($Cl^-$) | CATIONS ($Na^+, K^+$) | CATIONS ($Na^+, K^+$) |
| 6 | ***....hence...TYPE of st-st***<br>***ion homeostasis*** | ( [big $P_{Cl}$][small $I_{Naleak}$] )<br>**Donnan dominated** | ( [small $P_{Cl}$][big $I_{Naleak}$] )<br>**Pump-Leak Dominated** | ( [small $P_{Cl}$][big $I_{Naleak}$] )<br>**Pump-Leak dominated** |
| 7 | ***model complexity*** | **MINIMAL** P-L D system | minimal<br>(like SM-CD) | **NON-minimal** P-L D<br>(has K/Cl co-transporter)** |
| 8 | ***$P_{Cl}$ (molecular identity)*** | mostly ClC-1 | ---- | unclear (Rungta 2015) |
| 9 | ***$P_K$ (molecular identity)*** | girkS (DiFranco et al 2015),<br>K-ATP, others (needs work) | ---- | diverse, including 2 pore<br>domain $K^+$ channels |
| 10 | ***$P_{Na}$ (molecular identity)*** | **UNKNOWN** | ---- | NALCN, in part |

\* The teleost bodyform is broadly similar to that of the extinct lobefin fishes close to the common ancestors of mammals and teleosts (see Casane and Laurenti, 2013) and so, finding no estimates for ancestral vertebrates, %-ranges here for them are conservatively gauged from data for extant adult teleosts (typically ~60% and ~0.15% of total body mass for skeletal muscle (Johnston et al 2011; Helfman et al 2009) and an extant lobefin, whose brain occupies <<0.15% of body mass (Dutel et al 2019).
\*\*The K/Cl cotransporter of CN-CD is eliminated in MN-CD (Minimal Neuron"-CD) for which $V_{rest} = E_{Cl}$ ; **Table 2** gives parameters and steady-state values for both. Steady-state comparisons show: the cotransporter strongly hyperpolarizes $E_{Cl}$ , reduces $Vol_{cell}$ and minimally affects $[K^+]_i$ , $[Na^+]_i$ , $V_{rest}$ , ATP-consumption (as appropriate, the impacts of adding (e.g.) various $Na^+$ -transporters to SM-CD could similarly be probed).

### Ion homeostasis modeling via the charge difference approach

To explain the self-restoring V_rest_ of frog SMFs and the Na^+^-transporter regulation of SMF volume, Fraser and Huang (2004) introduced a charge difference (CD) approach. It explicitly includes Donnan effectors and counts all transmembrane ion and H_2_O movements. This enables simultaneous computation of steady-state Vol_cell_, [ion]_i_ and V_m_ (via charge on C_m_) (Fraser and Huang 2007, Usher-smith et al 2009; Kay 2018; Dmitriev et al 2019). Where relevant issues warrant it, CD-model trajectories of perturbed (non-steady-state) systems can be examined on timescales from milliseconds to days. Fraser and Huang’s (2004) seminal work described non-excitable SMFs. A later elaboration addresses APs and ion homeostasis in t-tubules (Fraser et al 2011). SM-CD, the excitable SMF model devised is as simple as an autonomous ion homeostasis model can be. It depicts not an entire multinucleate SMF but, to enable direct comparison with CN-CD, an existing (Dijkstra et al 2016) CD model for cortical neurons (**Figure 8 A,B,C**) an iterable fraction of a SMF (**Tables 1** and **2, Figure 8C,D**).

**FIGURE 2.**
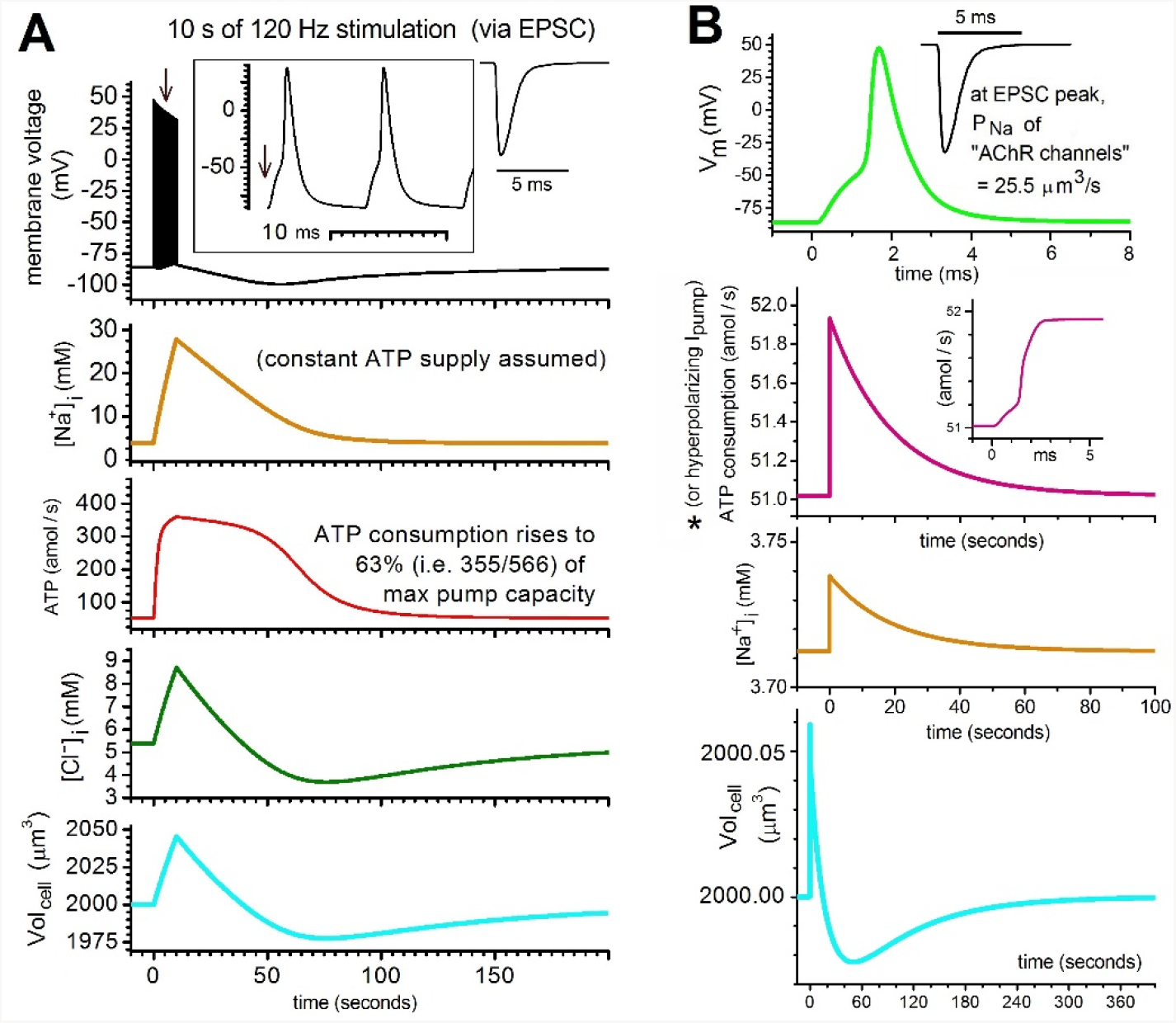
The P-L/D model, SM-CD, handling a real- life ion homeostatic stress-test. **A**. SM-CD undertaking a stress-test based on a rat soleus muscle experiment (Nielsen and Clausen 1997). SM-CD parameters are in **Tables 2**. H-H properties of the Nav and Kv channels are as depicted in **Figure 9C**, with rate constants as in **Table 4**. AP heights decline during the train. [Na^+^]_i_ rises, eliciting ↑ATP-consumption, ↑[Cl^-^]_i_, ↑Vol_cell_ and (not shown) ↓[K^+^]_i_. The expanded inset shows APs at ∼5 seconds. When EPSCs stop, V_m_ hyperpolarizes and [Na^+^]_i_ decays steadily due to electrogenic Na^+^-extrusion (hyperpolarizing I_pump_). ATP-consumption, having ↑∼7-fold (engaging ∼2/3 of SM-CD’s ∼11-fold pump-reserve) remains high for ∼40 seconds (“11-fold” refers to the maximal/resting ratio for ATP-consumption (i.e., 566/51.1; see **Figure 9A1**). Na^+^-extrusion, (I_pump_(t)) hyperpolarizes V_m_ until [Na^+^]_i_ returns to steady-state (**Table 2** gives models’ steady-state values). During the AP train, SM-CD steadily swells though by <2.5%; depending on a SMF’s geometry and mechanics, this might engage its caveolar tension-buffer (see Sinha et al 2011.; Morris 2018), a system not included in SM-CD. Once AChR, Nav, and Kv channel activity ceases, fluxes are strictly ion homeostatic. For ∼2 min, [Cl^-^]_i_ and Vol_cell_ undershoot, then converge on steady-state. Though ΔVol_cell_ is small [Cl^-^]_i_ almost doubles; new techniques (DiFranco et al 2019) would make this measurable. SM-CD’s very small P_Na_ ensures that the transiently-large I_pump_ is not undermined by a concurrently large I_Naleak_. The passive ion homeostatic processes, obeying the constraints set by Donnan effector electrostatics/thermodynamics (rapidly, thanks to SM-CD’s [big P_Cl_]), entrain to the [Na^+^]_i_-dependent Na^+^-pumping. H_2_O permeability is high enough for swelling to be effectively instantaneous; very small driving forces on Cl^-^ explain why (in spite of [big P_Cl_]) [Cl^-^]_i_ is slowest to converge back to steady-state. Vol_cell_(t) mirrors [Cl^-^]_i_(t). It would lag volume only if H_2_O permeability was rate-limiting. **B**. For a single AP, precisely the same phenomena are seen; notice that the time scale for Vol_cell_ is the longest (not shown in this case, but identical to [Cl^-^]_i_(t)). Notice the reminder (*) that ATP-consumption is directly proportional to (hyperpolarizing) I_pump_ (paired plots, **Figure 9A**). **Figure 6** provides further biophysical details.

**FIGURE 3.**
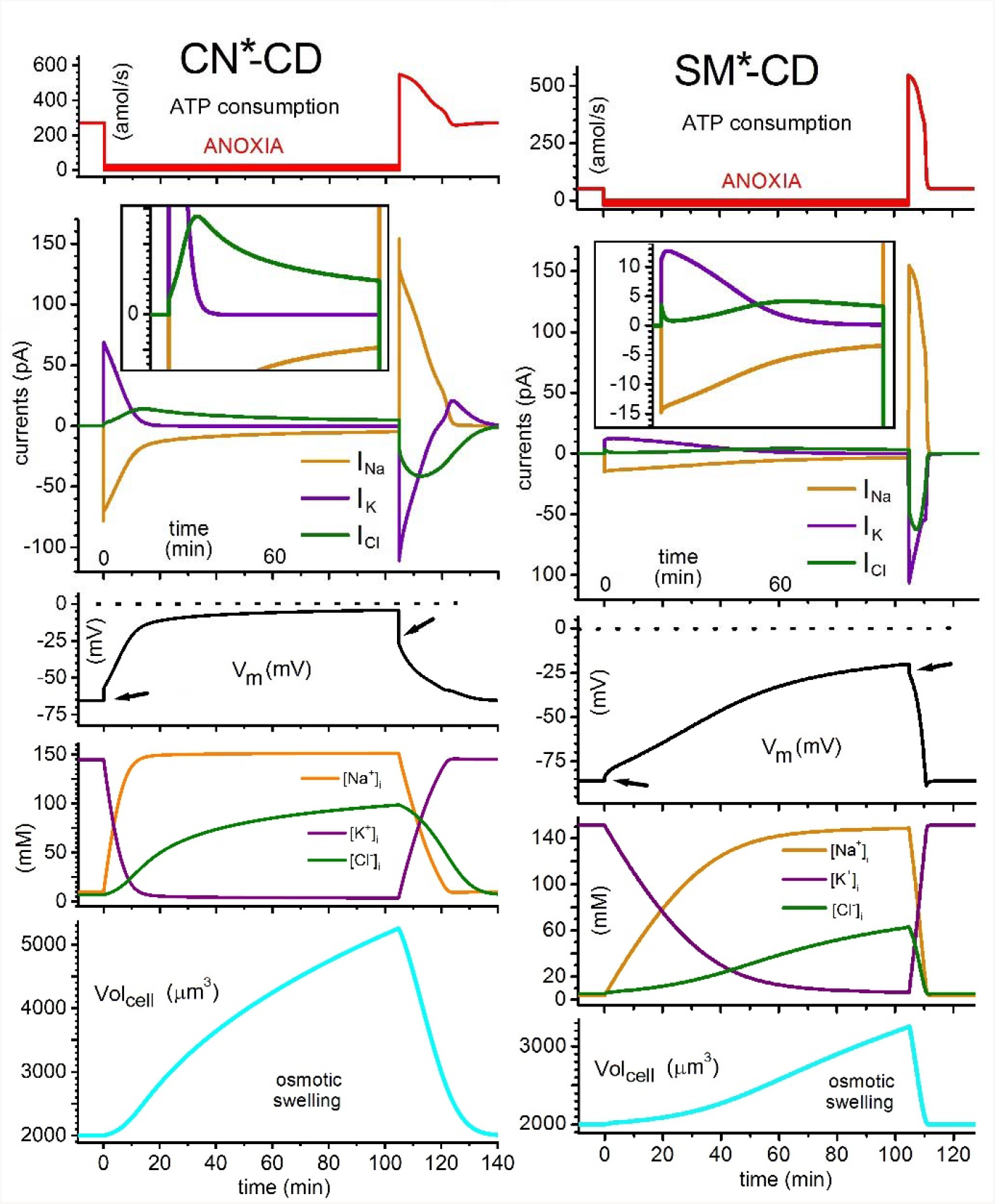
Anoxic rundown in inexcitable neuron and SMF models. **A**. CN*-CD and **B**. SM*-CD are CN-CD and SM-CD with V-gated channels (gNav, gKv plus “gCl(V)” in CN-CD) zeroed. Systems (non-minimal CN*-CD, minimal SM*-CD) start at steady-state. In V_m_ panels, arrows highlight the ΔV_m_ at I_pump_-off and I_pump_-on. Pump-off depolarizes CN*-CD by ∼8 mV, SM*-CD by <2 mV (for excitable CN-CD, the pump-off ΔV_m_ triggers APs; see **Figure 4A**). The SM*-CD rundown resembles Fig. 2 of Fraser and Huang (2004) (inexcitable amphibian SMF, V_rest_=-90 mV, more complex pump formulation). At t=0, driving forces are small on Cl^-^ and K^+^, large on Na^+^. Current trajectories show fluxes through P_Na_ and P_K_ (both bigger-valued in CN*-CD than in SM*-CD) dominating early rundown (though initially K^+^ feels the smaller driving force, I_Naleak_ through P_Na_ is the limiting cation flux). Swelling (Vol_cell_ tracks [Cl^-^]_i_) is checked far more effectively by SM*-CD’s extremely small P_Na_ than by CN*-CD’s [small P_Cl_]; CN*-CD swells 1.4X in 20 min, while SM*-CD swells 1.4X in 76 min. At 105 min (at pump-on) [Na^+^]_i_ maximally activates (566 amol/s) pumps in CN*-CD. To reach the same loss of gradient as CN*-CD at 105 min, anoxic rundown in SM*-CD would need to be extended to 15.5 hrs. Mainly because of its greater rundown at 105 min, recovery is slower in CN*-CD. Both models also recover fully at pump-on when anoxia is imposed for a total of 272 minutes (not shown). If recovery had been started at the same membrane voltage in both systems, CN*-CD would still recover more slowly because of its larger P_Na_.

**FIGURE 4.**
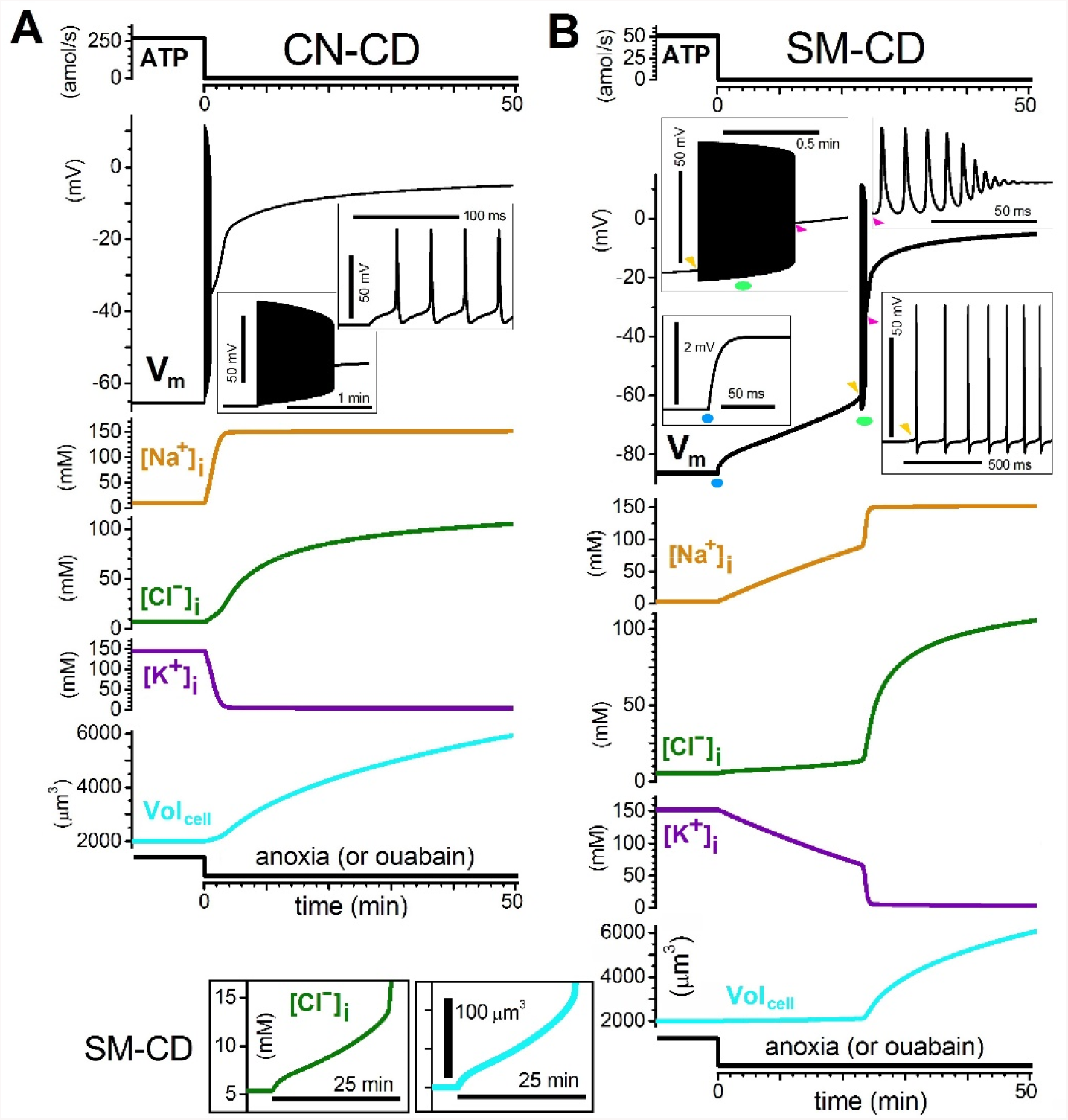
Anoxic rundown of CN-CD and SM-CD. V_m_(t) panels in **A** and **B** include expanded insets; the main trace in **B** (SM-CD) flags several insets. The blue dot expansion box shows that SM-CD switch-off of hyperpolarizing I_pump_ causes a depolarizing “RC” response in contrast to CN-CD’s larger pump-off depolarization which (**A**, inset) triggers ectopic APs. The 50 mV scale, lower right inset, **B**, applies also for the upper right inset. **A)** switch-off of CN-CD’s large steady-state hyperpolarizing I_pump_ triggers ectopic APs; the pump-off depolarization (readily evident in CN*-CD, **Figure 3A**) triggers ectopic APs that continue for <1 minute. Nav and Kv channels precipitously dissipate the cation gradients. This already-dire situation is exacerbated by the (pathological) opening at ∼-20 mV, of a depolarization-activated Cl^-^ pathway (Rungta at a 2015; Dijkstra et al 2016). CN-CD continues swelling, degrading towards DE. **B)** In SM-CD, I_pump_ switch-off produces a ∼2 mV RC response (as in **Figure 3**, where this is too fast to be temporally-resolved) then a very slow V_m_ rundown whose rate is limited by small P_Na_. [Big P_Cl_] lets E_Cl_ rapidly track V_m_. Because P_K_>>P_Na_, K^+^ efflux mostly neutralizes Na^+^ influx. Slight swelling reflects the extent of [Na^+^+Cl^-^]_i_ entry, osmo-balanced by H_2_O influx (but the Cl^-^ driving force is negligible). Ectopic firing starts at 22.7 minutes. SM-CD Nav and Kv density is 3X greater than in CN-CD, so gradient dissipation is even more precipitous. As E_K_ decreases during APs, Na^+^ influx is neutralized increasingly by Cl^-^ influx, and since the [Na^+^+Cl^-^]influx→[Na^+^+Cl^-^+H_2_O]influx, Vol_cell_ suddenly increases. Ectopic firing tapers abruptly, the small V_m_ oscillations reflecting V-gating (SM-CD, pink triangle, V_m_ inset). When [Na^+^]_i_ and [K^+^]_I_ attain external values, V_m_ is still <0 mV and [Cl^-^]_I_<[Cl^-^]_e_ so net Cl^-^ entry continues, accompanied by H_2_O for osmo-balance; compartment neutrality is retained during swelling (ongoing ↑H_2_O) because ↑[Cl^-^]_i_ compensates for ↓[impermanent anion]_i_. The system would rupture before converging on DE (see **Table 1**).

**FIGURE 5.**
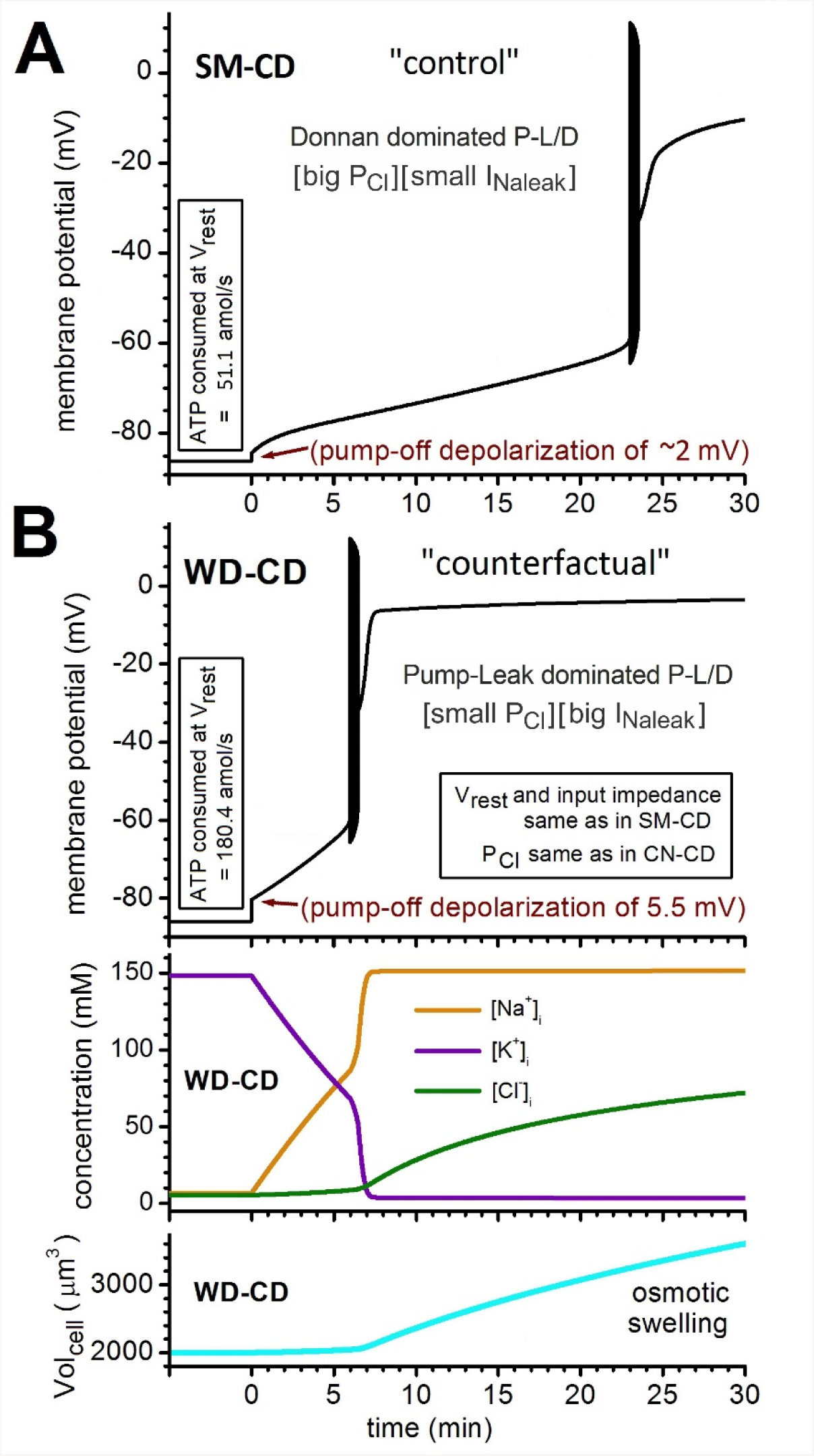
WD-CD, a counterfactual analog of SM-CD. To clarify how [big P_Cl_] plays out in the context of SM-CD’s other ion homeostatic elements (beyond its electrophysiological role of “big input conductance”), anoxic rundown is compared in WD-CD, a minimal P-L/D model with big P_K_ as “big input conductance”. **A** (“control”) is the SM-CD anoxic rundown of V_m_ (for other parameters, see **Figure 4B**). **B**, as labeled is the WD-CD anoxic rundown. WD-CD’s ion homeostatic response to an EPSC-triggered AP is shown in **Figure 6**. WD-CD was subjected to the **Figure 2A** stress test (see **Figure Supp01C**).

**FIGURE 6.**
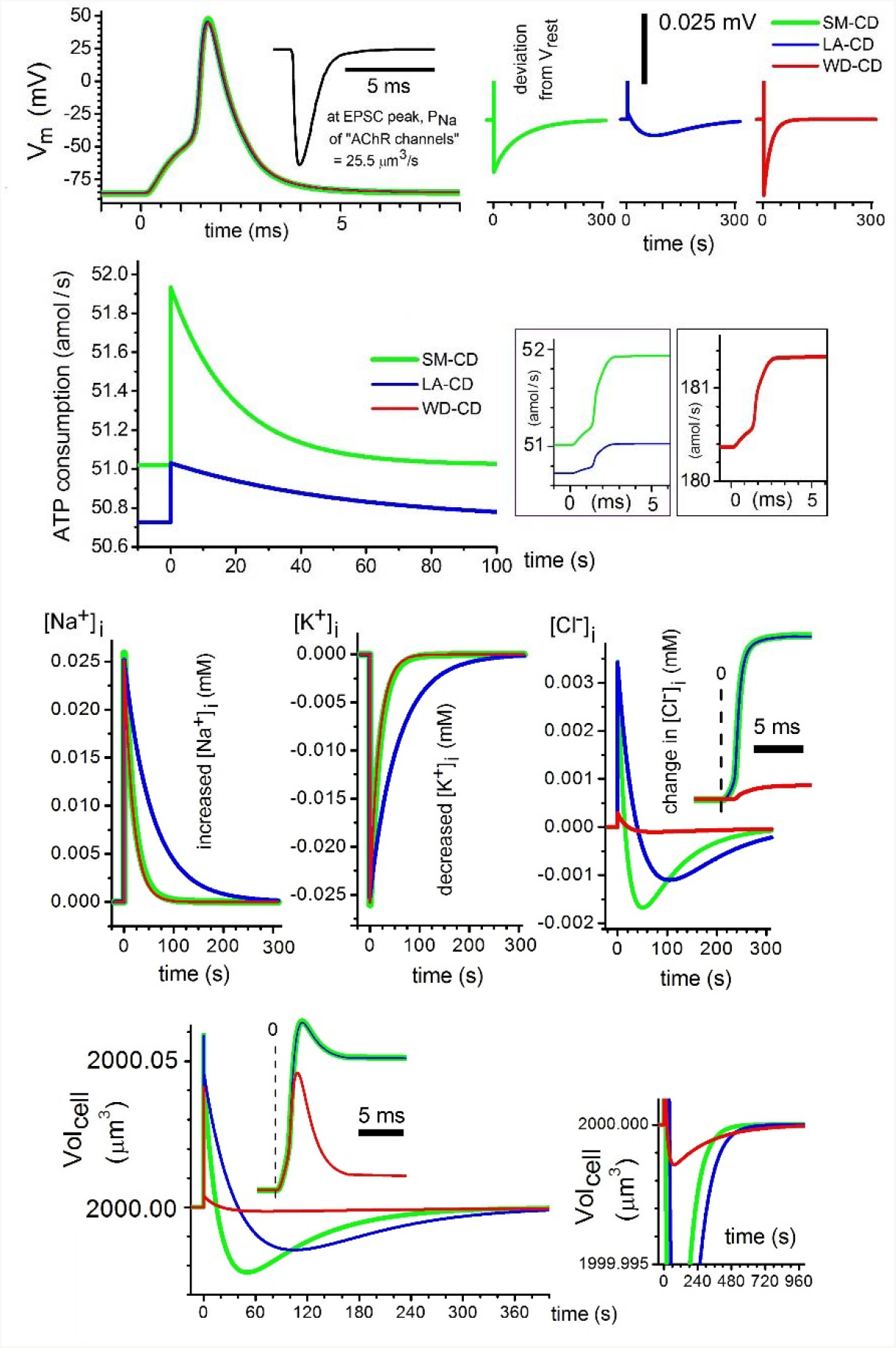
Biophysics of Donnan dominated and Pump-Leak dominated systems. Ion homeostasis is scrutinized comparatively for 3 related minimal P-L/D systems: they are the normal version and an ischemic version (LA-CD; 30% pump-strength) of SM-CD, both of which are Donnan dominated systems, and SM-CD’s Pump-Leak dominated counterfactual analog, WD-CD. In addition to the APs and subsequent ΔV_m_, ATP-consumption (directly proportional to I_pump_), Δ[ion]_i_ for (Na^+^, K^+^ and Cl^-^) and Vol_cell_ are plotted. In each system, ion homeostatic restoration to steady-state involves a volume oscillation commencing with AP-induced “H_2_O blips” which are abrupt ↑Vol_cell_ (grey arrow) whose amplitude reflects the magnitude of excess Na^+^ entry (i.e., the Na^+^-entry demanding osmo-balance because it is unaccompanied by K^+^-exit). This excess is greater in SM-CD and LA-CD (whose blips overlap) than in WD-CD (whose bigger P_K_ explains its smaller blip). The amplitude of the models’ immediate Donnan-responses (see millisecond resolution trajectories for the ↑[Cl^-^]_i_(t)) depend on what fraction of the “electro-neutralization-of-excess-Na^+^-entry” task is achieved by Cl^-^ entry (for [big P_Cl_] SM-CD and LA-CD, most of it; for [small P_Cl_] WD-CD, very little of it). The AP perturbation ends in milliseconds but ion homeostatic feedback continues for minutes, the pump sensor/effector process actively extruding Na^+^ (=hyperpolarizing I_pump_) at a rate proportional to [Na^+^]_i,_ while the passive Donnan sensor/effector feedback mechanism supports Cl^-^ and H_2_O fluxes that keep in/out H_2_O activities balanced and the cytoplasm electroneutral.

**FIGURE 7.**
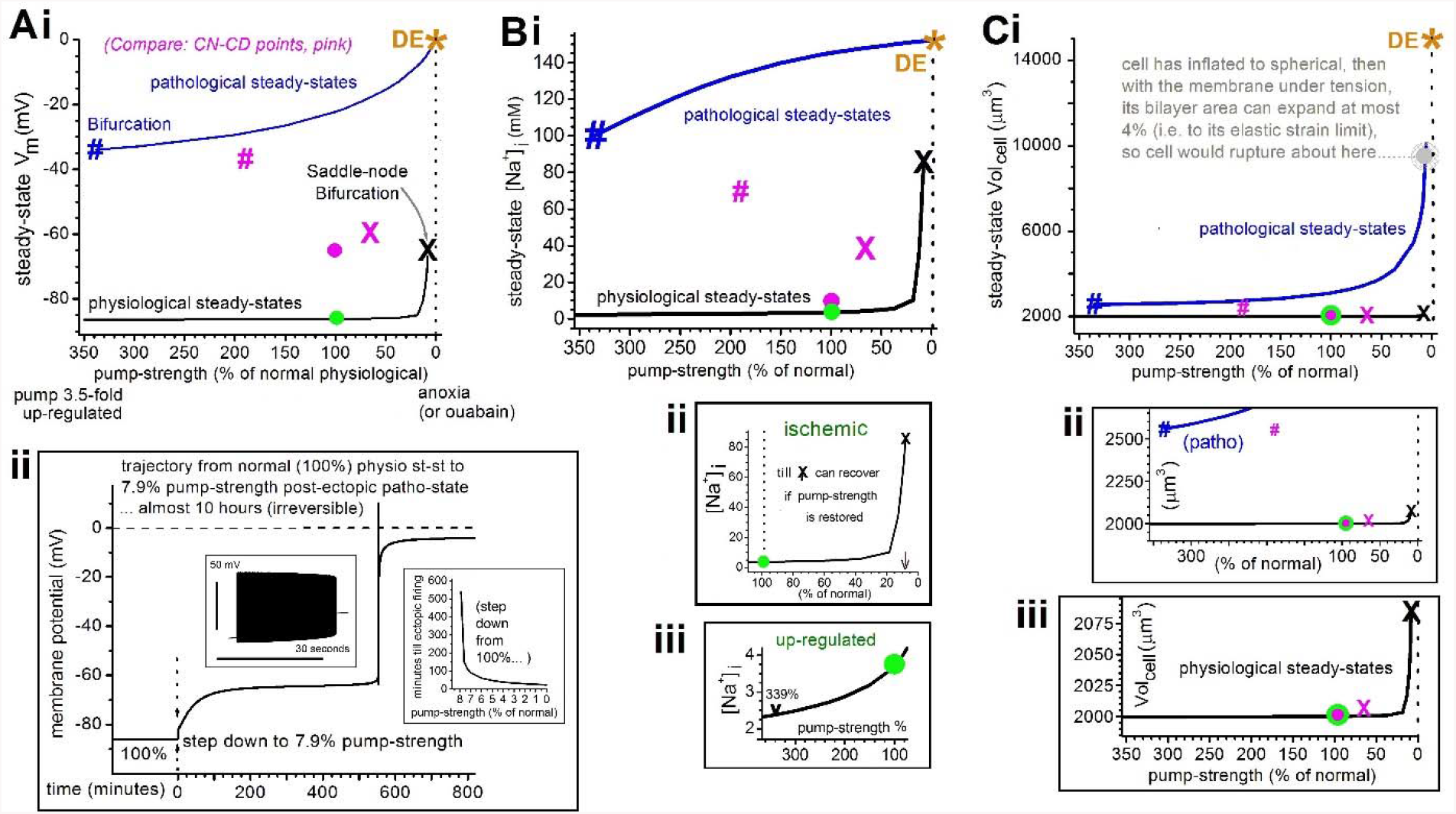
SM-CD bistability: bifurcation plots. **Ai,Bi,Ci** are SM-CD bifurcation plots, as labeled (**Figure Supp 02A,B** has [K^+^]_i_ and [Cl^-^]_i,_ bifurcation plots). Exit from the physiological continuum occurs at the “saddle-node” marked **X**; exit from the pathological continuum occurs at the unstable point marked **#**. Parts of **Bi** and **Ci** are expanded (**Bii,iii** and **Cii,iii**). **Aii** is the V_m_(t) trajectory for 100%→7.9% pump-strength (i.e. saddle-node); it takes ∼10 hours for SM-CD to depolarize to firing threshold (inset expands ectopic burst) and then re-stabilize in a pathological steady-state. For bigger pump-strength drops, time-to-ectopic-firing falls sharply (inset plot), converging on the anoxic (0%) rundown time (22.7 minutes) (see **Figure 4A**). For patho-physiological comparisons, we recomputed the CN-CD (Dijkstra et al 2016) bifurcation plots (we report Vol_cell_, not cross-sectional area); CN-CD’s threshold and normal steady-state points are included, as indicated. Although notional Donnan Equilibrium (DE) is marked in **Ai,Bi,Ci**, rupture would occur prior to attaining DE (as per **Ci**) unless bilayer area was augmented (↑C_m_).

**FIGURE 8.**
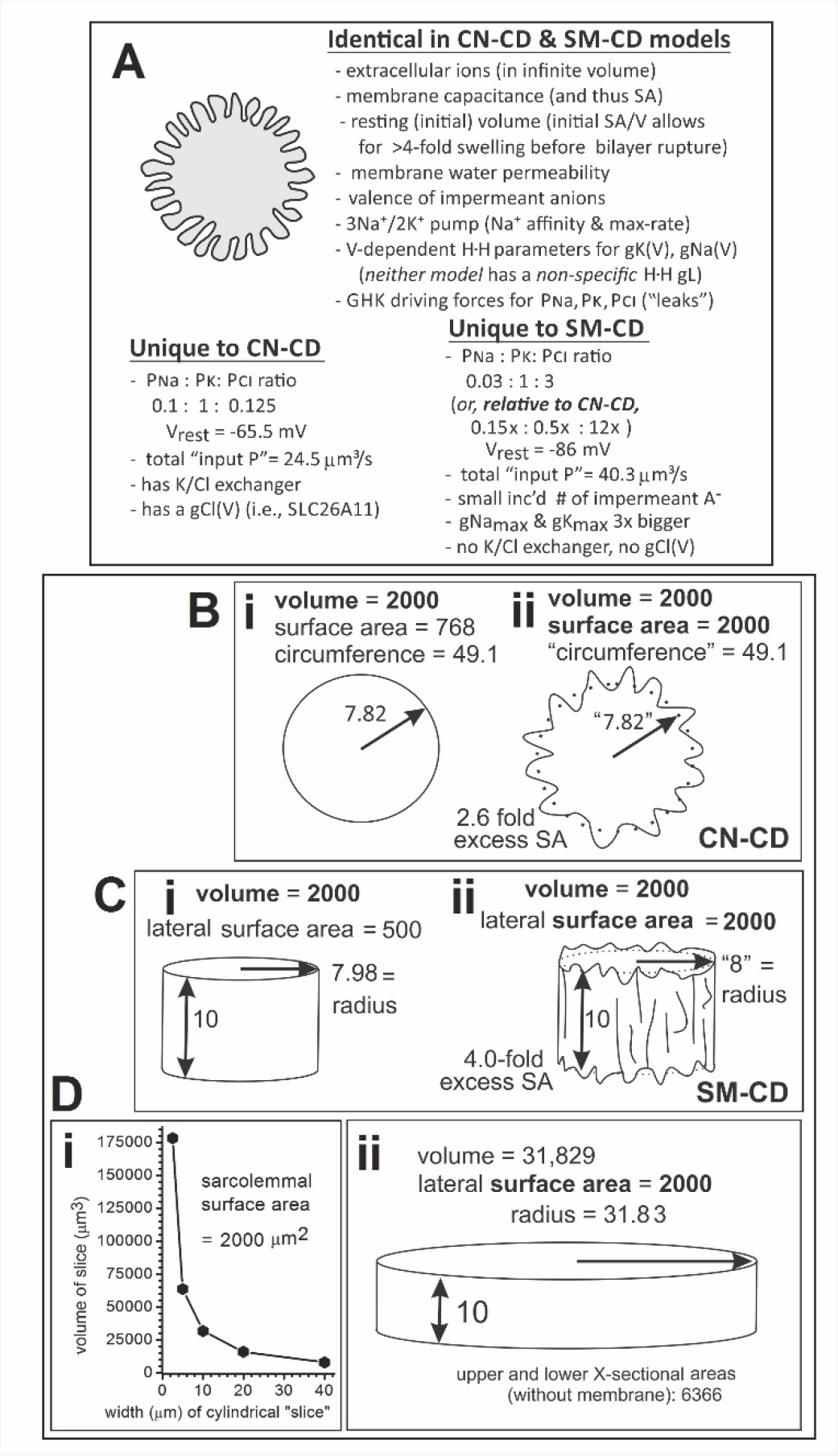
Model building. **A**. **From CN-CD to SM-CD** SM-CD was devised for direct comparison against CN-CD, an established cortical neuron model (Dijkstra et al 2016). Where possible the two are identical. Differences reflect the distinctive electrophysiological “lifestyles” of neurons (Dijkstra et al 2016) and SMFs (Fraser and Huang, 2004; Fu et al 2011; Ruff 2011; Pedersen et al 2011; Novak et al 2015). The Dijkstra et al (2016) model addressed cortical neurons in brainslice preparations. As an indicator of Vol_cell_ they reported confocally-imaged cross-sectional neuronal areas; we recalculated to obtain spherical volumes from those X-areas. **B**,**C**,**D**. **From CN-CD to SM-CD to SMF-like cytomorphology** To foster comparisons, Vol_cell_=2000 µm^3^ and SA=2000 µm^2^ from CN-CD (Dijkstra et al 2016) was used for SM-CD so there is excess SA beyond what is required to enclose a sphere of 2000 µm^3^, but neuron and SMF cytomorphologies differ, so **Bi,ii**, and **Ci,ii** sketches the disposition of the excess SA (as per **A**) for each. Additionally, **C** indicates that for SM-CD, the 2000 µm^3^ Vol_cell_ would represent one cross-sectional slice “ion homeostatic unit” through a cylindrical multinucleate myofiber. The 2000 µm^2^ of sarcolemma would encircle, not enclose, myoplasm. **Cii** is thus “SM-CD” as modeled here (Vol_cell_=2000 µm^3^, SA =2000 µm^2^). The P-L/D models are not, however, 3-dimensional, so the difference noted in the excess SA for CN-CD and SM-CD does not affect computations here. However, as **Di** shows, for cylindrical geometry, 2000 µm^2^ of sarcolemma disposed as a smooth (no excess-SA) ring of SA could encircle vastly more than Vol_cell_ =2000 µm^3^, depending on slice width. For example, the 10 µm slice depicted in **Dii** would enclose almost 32,000 µm^3^ of myoplasm. Or, for an example with a little excess SA, 2000 µm^2^ could be disposed, say, as follows (not shown): a 15 µm radius cylinder of 20 µm “slice width” with a lateral SA=1885 µm^2^ encircling 14,137 µm^3^ with 115 µm^2^ added to bring the slice to SA=2000 µm^2^ and providing a small (1.06-fold) tension-buffering excess SA. Throughout the figure, linear, area and volume dimensions have units of µm, µm^2^ and µm^3^ respectively. Schematics are not to scale. For reference, Fraser and Huang (2004) modeled a 75 µm diameter fiber with SA/Vol_cell_ =5×10^5^ cm^2^/liter; the **Dii** slice has SA/Vol_cell_ =6×10^5^ cm^2^/liter (for SM-CD as per **Cii**, SA/Vol_cell_ =25×10^5^ cm^2^/liter).

CD modeling comparisons here highlight two overlooked facets of SMF physiology: **1)** the collaboration of ClC-1 channels with Donnan effectors is imperative for SMF ion homeostasis (not just for low SMF excitability (Jentsch and Pusch, 2018; Jeng et al 2020) and **2)** the extraordinary smallness of the (unidentified) SMF P_Na_ renders it pivotal (via its [small I_Naleak_]) to the robustness of SMF ion homeostasis.

### Donnan forces finessed as “found free-energy” for volume homeostasis

Though ion homeostasis is commonly referred to as a Pump-Leak process, dropping “Donnan” abrogates the P-L/D process; the Donnan-mediated volume feature is mandatory. **Figure 1B** (and legend) introduces the simple but novel idea that at steady-state, P-L/D ion homeostasis can be Donnan dominated or Pump-Leak dominated. Each of the two systems depicted has the minimal components required for autonomous P-L/D ion homeostasis; they differ in relative densities of always-open (“resting” or “leak”) channels, P_Cl_, P_K_ and P_Na_.

At steady-state in the Donnan dominated system, the transmembrane flux of ions is due mostly to Cl^-^ ions, with influx and efflux perfectly balanced thanks to the cytoplasm’s impermeant anions, while at steady-state in the Pump-Leak dominated system, the transmembrane ion flux is due mostly to cations, with passive Na^+^-influx and K^+^-efflux perfectly balanced by active (pumped) counterfluxes. Animal cells’ ever-present Donnan forces are seen as a potentially lethal problem for cells to solve. When pumps falter, after all, [Na^+^+Cl^-^+H_2_O] entry kills cells as a direct consequence of Donnan forces. Animal cells handle the [Na^+^+Cl^-^+H_2_O]-danger by sequestering the perilous free energy. Neurons use [small P_Cl_], SMFs use [small P_Na_].

Neuronal steady-state is thus a [small P_Cl_][big I_Naleak_] Pump-Leak dominated process while SMF steady-state is a [big P_Cl_][small I_Naleak_] Donnan dominated process. Till now however, the distinction between Donnan dominated and Pump-Leak dominated strategies has, however, gone unrecognized (or at least unnamed). ClC-1 channels, expressed abundantly and exclusively in SMFs, are the major component (Pedersen et al 2016) of their [big P_Cl_]. Discovered in 1991 by the Jentsch lab (Steinmeyer et al 1991) via close homology to elasmobranch ClC-0 (see Miller 2015), their role is described in electrophysiological terms; i.e, ClC-1 passively stabilizes the hyperpolarized V_rest_ (Jentsch & Pusch 2018). The P-L/D framework shows that, simultaneously, ClC-1 is crucial for SMF ion homeostasis. A counterfactual model (a big P_K_ SMF-analog P-L/D) helps clarify this point.

Cytoplasmic proteins did not evolve to serve ion homeostasis. Nevertheless the unavoidable “colligative-charge” attribute that makes them Donnan effectors is a source of “found free energy”, one that cells use in a sensor/effector role for ion homeostatic feedback (**Figure 1A**). Low-excitability SMFs evolved finessing their Donnan effector/sensor system, making it a high-conductance (=low-impedance) stabilizer of their low-cost ion homeostatic steady-state (E_Cl_=V_rest_). Modeling here reveals how, by maintaining [small I_Naleak_] and exploiting that found free energy, SMFs obtain superb operational resilience. Donnan dominated SMF ion homeostasis, we propose, played an important role in the vertebrate lineage. Our initial motivation was, however, to understand SMF longevity in Duchenne muscular dystrophy (DMD) patients (Allen et al 2016; Gerhalter et al 2019) where SMFs experience chronic ion homeostatic stresses that rapidly kill neurons. In a companion paper (Morris et al submitted), SM-CD, the P-L/D model for SMF ion homeostasis explained here, is used to address that issue.

## RESULTS

### SM-CD passes a stress-test

How SM-CD was devised is explained in **Methods**. Because SM-CD is new, we begin by demonstrating its general merit via a “reality check” (**Figure 2A**). Clausen (2015) summarized experimental findings (Nielsen and Clausen 1997) as follows: “In isolated rat soleus… after 10 sec of stimulation at 120 Hz, the net Na^+^ re-extrusion measured in the subsequent 30 sec of rest reaches a 22-fold increase in Na^+^, K^+^ pump activity corresponding to 97% of the theoretical maximum rate of active Na^+^, K^+^ pumping measured and calculated on the basis of total content of ^3^H-ouabain binding sites”. **Figure 2A** follows V_m_, [Na^+^]_i_, [Cl^-^]_i,_ Vol_cell_ and ATP-consumption in SM-CD while 1200 APs fire and the while SM-CD fully restores itself to steady-state. The “Na^+^ re-extrusion” time course is reassuringly similar to rat soleus. In SM-CD, the maximal ↑I_pump_ is 11.1-fold. When increased to 22.2-fold (**Figure Sup1A)** ATP-consumption(t) responds accordingly. **Figure 2B** shows one EPSC-elicited AP in SM-CD and the trajectories (milliseconds, minutes) of its ion homeostatic parameters. Having demonstrated SM-CD’s response to challenge, we compare neuron versus SMF P-L/D processes. The biophysics of single AP ion homeostasis will be addressed later (**Figure 6**).

### Steady-state, anoxic rundown, recovery: P-L/D systems without excitability

If their sole energy-generator (hyperpolarizing I_pump_) is eliminated, P-L/D systems degrade towards Donnan equilibrium (DE). Dijkstra et al (2016) mimicked brain-slice anoxia in CN-CD, making switch-off gradual (like wash-in of pump-inhibitors). Here, for biophysical clarity, “pump-off/on” is instantaneous.

**Table 2.**
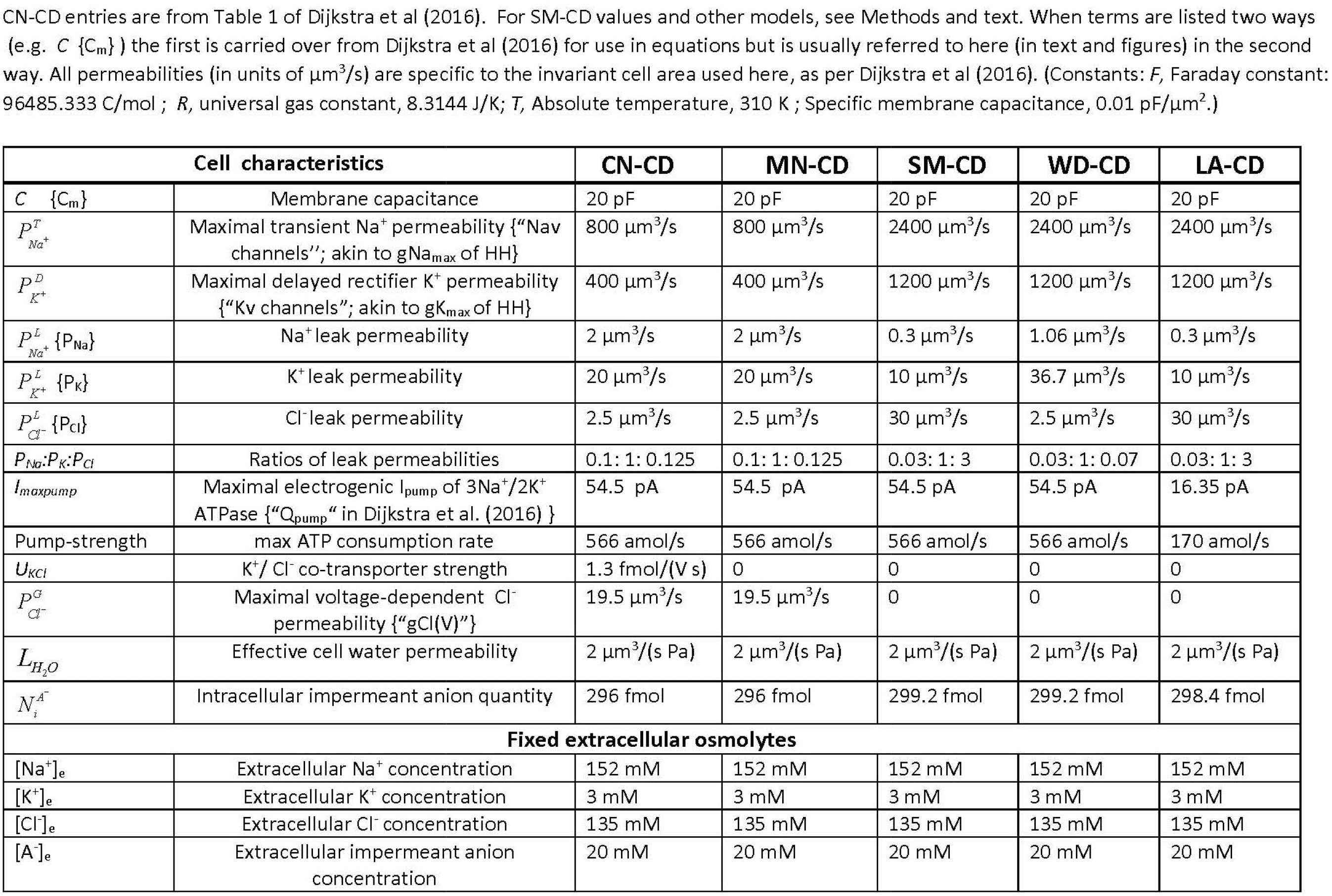
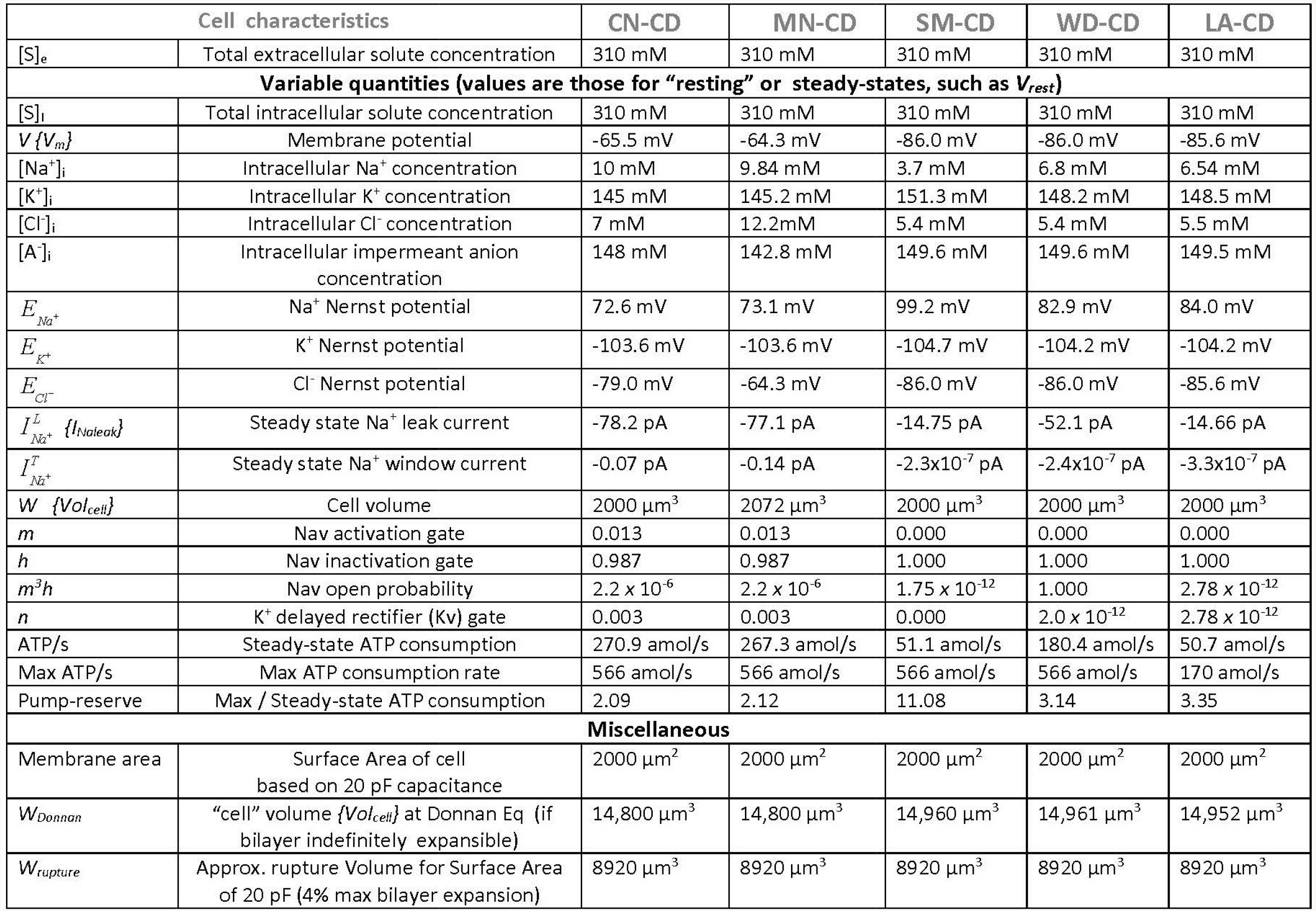
Parameter values for CN-CD and SM-CD models and variants.

| Cell characteristics |  | CN-CD | MN-CD | SM-CD | WD-CD | LA-CD |
| --- | --- | --- | --- | --- | --- | --- |
| $C \{C_m\}$ | Membrane capacitance | 20 pF | 20 pF | 20 pF | 20 pF | 20 pF |
| $P_{Na^+}^T$ | Maximal transient $\text{Na}^+$ permeability {"Nav channels"; akin to $g_{\text{Na}_{\text{max}}}$ of HH} | 800 $\mu\text{m}^3/\text{s}$ | 800 $\mu\text{m}^3/\text{s}$ | 2400 $\mu\text{m}^3/\text{s}$ | 2400 $\mu\text{m}^3/\text{s}$ | 2400 $\mu\text{m}^3/\text{s}$ |
| $P_{K^+}^D$ | Maximal delayed rectifier $\text{K}^+$ permeability {"Kv channels"; akin to $g_{\text{K}_{\text{max}}}$ of HH} | 400 $\mu\text{m}^3/\text{s}$ | 400 $\mu\text{m}^3/\text{s}$ | 1200 $\mu\text{m}^3/\text{s}$ | 1200 $\mu\text{m}^3/\text{s}$ | 1200 $\mu\text{m}^3/\text{s}$ |
| $P_{Na^+}^L \{P_{Na}\}$ | $\text{Na}^+$ leak permeability | 2 $\mu\text{m}^3/\text{s}$ | 2 $\mu\text{m}^3/\text{s}$ | 0.3 $\mu\text{m}^3/\text{s}$ | 1.06 $\mu\text{m}^3/\text{s}$ | 0.3 $\mu\text{m}^3/\text{s}$ |
| $P_{K^+}^L \{P_K\}$ | $\text{K}^+$ leak permeability | 20 $\mu\text{m}^3/\text{s}$ | 20 $\mu\text{m}^3/\text{s}$ | 10 $\mu\text{m}^3/\text{s}$ | 36.7 $\mu\text{m}^3/\text{s}$ | 10 $\mu\text{m}^3/\text{s}$ |
| $P_{Cl^-}^L \{P_{Cl}\}$ | $\text{Cl}^-$ leak permeability | 2.5 $\mu\text{m}^3/\text{s}$ | 2.5 $\mu\text{m}^3/\text{s}$ | 30 $\mu\text{m}^3/\text{s}$ | 2.5 $\mu\text{m}^3/\text{s}$ | 30 $\mu\text{m}^3/\text{s}$ |
| $P_{Na^+}P_{K^+}P_{Cl^-}$ | Ratios of leak permeabilities | 0.1 : 1 : 0.125 | 0.1 : 1 : 0.125 | 0.03 : 1 : 3 | 0.03 : 1 : 0.07 | 0.03 : 1 : 3 |
| $I_{\text{max pump}}$ | Maximal electrogenic $I_{\text{pump}}$ of $3\text{Na}^+/2\text{K}^+$ ATPase {"$Q_{\text{pump}}$" in Dijkstra et al. (2016) } | 54.5 pA | 54.5 pA | 54.5 pA | 54.5 pA | 16.35 pA |
| Pump-strength | max ATP consumption rate | 566 amol/s | 566 amol/s | 566 amol/s | 566 amol/s | 170 amol/s |
| $U_{KCl}$ | $\text{K}^+/\text{Cl}^-$ co-transporter strength | 1.3 fmol/(V s) | 0 | 0 | 0 | 0 |
| $P_{Cl^-}^G$ | Maximal voltage-dependent $\text{Cl}^-$ permeability {"$g_{\text{Cl(V)}}$"} | 19.5 $\mu\text{m}^3/\text{s}$ | 19.5 $\mu\text{m}^3/\text{s}$ | 0 | 0 | 0 |
| $L_{H_2O}$ | Effective cell water permeability | 2 $\mu\text{m}^3/(\text{s Pa})$ | 2 $\mu\text{m}^3/(\text{s Pa})$ | 2 $\mu\text{m}^3/(\text{s Pa})$ | 2 $\mu\text{m}^3/(\text{s Pa})$ | 2 $\mu\text{m}^3/(\text{s Pa})$ |
| $N_i^A$ | Intracellular impermeant anion quantity | 296 fmol | 296 fmol | 299.2 fmol | 299.2 fmol | 298.4 fmol |
| Fixed extracellular osmolytes |  |  |  |  |  |  |
| $[\text{Na}^+]_e$ | Extracellular $\text{Na}^+$ concentration | 152 mM | 152 mM | 152 mM | 152 mM | 152 mM |
| $[\text{K}^+]_e$ | Extracellular $\text{K}^+$ concentration | 3 mM | 3 mM | 3 mM | 3 mM | 3 mM |
| $[\text{Cl}^-]_e$ | Extracellular $\text{Cl}^-$ concentration | 135 mM | 135 mM | 135 mM | 135 mM | 135 mM |
| $[\text{A}^-]_e$ | Extracellular impermeant anion concentration | 20 mM | 20 mM | 20 mM | 20 mM | 20 mM |
| [S]<sub>e</sub> | Total extracellular solute concentration | 310 mM | 310 mM | 310 mM | 310 mM | 310 mM |
| Variable quantities (values are those for "resting" or steady-states, such as $V_{rest}$) | | | | | | |
| [S]<sub>i</sub> | Total intracellular solute concentration | 310 mM | 310 mM | 310 mM | 310 mM | 310 mM |
| $V\{V_m\}$ | Membrane potential | -65.5 mV | -64.3 mV | -86.0 mV | -86.0 mV | -85.6 mV |
| [Na<sup>+</sup>]<sub>i</sub> | Intracellular Na<sup>+</sup> concentration | 10 mM | 9.84 mM | 3.7 mM | 6.8 mM | 6.54 mM |
| [K<sup>+</sup>]<sub>i</sub> | Intracellular K<sup>+</sup> concentration | 145 mM | 145.2 mM | 151.3 mM | 148.2 mM | 148.5 mM |
| [Cl<sup>-</sup>]<sub>i</sub> | Intracellular Cl<sup>-</sup> concentration | 7 mM | 12.2 mM | 5.4 mM | 5.4 mM | 5.5 mM |
| [A<sup>-</sup>]<sub>i</sub> | Intracellular impermeant anion concentration | 148 mM | 142.8 mM | 149.6 mM | 149.6 mM | 149.5 mM |
| $E_{Na^+}$ | Na<sup>+</sup> Nernst potential | 72.6 mV | 73.1 mV | 99.2 mV | 82.9 mV | 84.0 mV |
| $E_{K^+}$ | K<sup>+</sup> Nernst potential | -103.6 mV | -103.6 mV | -104.7 mV | -104.2 mV | -104.2 mV |
| $E_{Cl^-}$ | Cl<sup>-</sup> Nernst potential | -79.0 mV | -64.3 mV | -86.0 mV | -86.0 mV | -85.6 mV |
| $I_{Na^+}^L \{I_{NaLeak}\}$ | Steady state Na<sup>+</sup> leak current | -78.2 pA | -77.1 pA | -14.75 pA | -52.1 pA | -14.66 pA |
| $I_{Na^+}^T$ | Steady state Na<sup>+</sup> window current | -0.07 pA | -0.14 pA | -2.3x10<sup>-7</sup> pA | -2.4x10<sup>-7</sup> pA | -3.3x10<sup>-7</sup> pA |
| $W\{Vol_{cell}\}$ | Cell volume | 2000 μm<sup>3</sup> | 2072 μm<sup>3</sup> | 2000 μm<sup>3</sup> | 2000 μm<sup>3</sup> | 2000 μm<sup>3</sup> |
| $m$ | Nav activation gate | 0.013 | 0.013 | 0.000 | 0.000 | 0.000 |
| $h$ | Nav inactivation gate | 0.987 | 0.987 | 1.000 | 1.000 | 1.000 |
| $m^3h$ | Nav open probability | 2.2 x 10<sup>-6</sup> | 2.2 x 10<sup>-6</sup> | 1.75 x 10<sup>-12</sup> | 1.000 | 2.78 x 10<sup>-12</sup> |
| $n$ | K<sup>+</sup> delayed rectifier (Kv) gate | 0.003 | 0.003 | 0.000 | 2.0 x 10<sup>-12</sup> | 2.78 x 10<sup>-12</sup> |
| ATP/s | Steady-state ATP consumption | 270.9 amol/s | 267.3 amol/s | 51.1 amol/s | 180.4 amol/s | 50.7 amol/s |
| Max ATP/s | Max ATP consumption rate | 566 amol/s | 566 amol/s | 566 amol/s | 566 amol/s | 170 amol/s |
| Pump-reserve | Max / Steady-state ATP consumption | 2.09 | 2.12 | 11.08 | 3.14 | 3.35 |
| Miscellaneous |  |  |  |  |  |  |
| Membrane area | Surface Area of cell based on 20 pF capacitance | 2000 μm<sup>2</sup> | 2000 μm<sup>2</sup> | 2000 μm<sup>2</sup> | 2000 μm<sup>2</sup> | 2000 μm<sup>2</sup> |
| $W_{Donnan}$ | "cell" volume {$Vol_{cell}$} at Donnan Eq (if bilayer indefinitely expansible) | 14,800 μm<sup>3</sup> | 14,800 μm<sup>3</sup> | 14,960 μm<sup>3</sup> | 14,961 μm<sup>3</sup> | 14,952 μm<sup>3</sup> |
| $W_{rupture}$ | Approx. rupture Volume for Surface Area of 20 pF (4% max bilayer expansion) | 8920 μm<sup>3</sup> | 8920 μm<sup>3</sup> | 8920 μm<sup>3</sup> | 8920 μm<sup>3</sup> | 8920 μm<sup>3</sup> |

**Table 4.** Voltage-dependent rates for H-H gating variables^*^

| Term | Expression | Description |
| --- | --- | --- |
| $\alpha_m(V)$ | $\frac{0.32(V + 52 \text{ mV})}{1 - \exp\left(-\frac{V + 52 \text{ mV}}{4 \text{ mV}}\right)} \frac{\text{kHz}}{\text{mV}}$ | Activation rate for transient Na<sup>+</sup> channel |
| $\beta_m(V)$ | $\frac{0.28(V + 25 \text{ mV})}{\exp\left(-\frac{V + 25 \text{ mV}}{5 \text{ mV}}\right) - 1} \frac{\text{kHz}}{\text{mV}}$ | Deactivation rate for transient Na<sup>+</sup> channel |
| $\alpha_h(V)$ | $0.128 \exp\left(-\frac{V + 53 \text{ mV}}{18 \text{ mV}}\right) \text{ kHz}$ | Inactivation rate for transient Na<sup>+</sup> channel |
| $\beta_h(V)$ | $\frac{4}{1 + \exp\left(-\frac{V + 30 \text{ mV}}{5 \text{ mV}}\right)} \text{ kHz}$ | Recovery from inactivation rate for transient Na<sup>+</sup> channel |
| $\alpha_n(V)$ | $\frac{0.016(V + 35 \text{ mV})}{1 - \exp\left(-\frac{V + 35 \text{ mV}}{5 \text{ mV}}\right)} \frac{\text{kHz}}{\text{mV}}$ | Activation rate for delayed rectifier K<sup>+</sup> channel |
| $\beta_n(V)$ | $0.25 \exp\left(-\frac{V + 50 \text{ mV}}{40 \text{ mV}}\right) \text{ kHz}$ | Deactivation rate for delayed rectifier K<sup>+</sup> channel |

\* Based on Kager et al (2000) but as used in Dijkstra et al (2016) with one typographical error corrected.

In **Figure 3**, to examine steady-state P-L/D components only, all V-gated channels are zeroed (*). For CN- or CN*-CD and for SM- or SM*-CD, steady-states are identical. Before t=0, currents=zero and Vol_cell_, V_rest_, [ion]_i_ and ATP-consumption are as per CN-CD, SM-CD, **Table 2**. At pump-off (t=0), rundown commences. Pumping is restored at 105 minutes (before swelling to rupture point, and long before “notional” DE; see **Table 2**, Miscellaneous).

Before t=0, steady-state ATP-consumption already tells a major story: CN*-CD maintains V_rest_=-65.5 mV for 270.9 amol/s, whereas SM*-CD maintain V_rest_=-86 mV for 51.1 amol/s.

The frugality of SM*-CD reflects the fact that ATP-consumption directly mirrors I_Naleak_ (as per **Figure 1B**) and SM*-CD is a [big P_Cl_][small I_Naleak_] system while CN*-CD is a [small P_Cl_][big I_Naleak_] system. CN-CD is non-minimal, but that negligibly affects its ATP-consumption (confirm by comparing CN-CD to MN-CD, **Table 2**). I_Naleak_ depends on P_Na_ and the electrodiffusive driving force (**Equations 2** and **3**). Though the Na^+^ driving force is 21 mV bigger in SM*-CD than CN*-CD at their respective V_rest_, this is outweighed by P_Na_ being 67X smaller in SM*-CD. Because I_pump_=–I_Naleak_, “instantaneous” (t=0) I_Na_ (**Figure 3A,B)** directly shows the crucial I_Naleak_ difference that makes steady-state ATP-consumption by SM(*)-CD <1/5.3 that of CN(*)-CD (see also **Figure 9Ai**).

**FIGURE 9.**
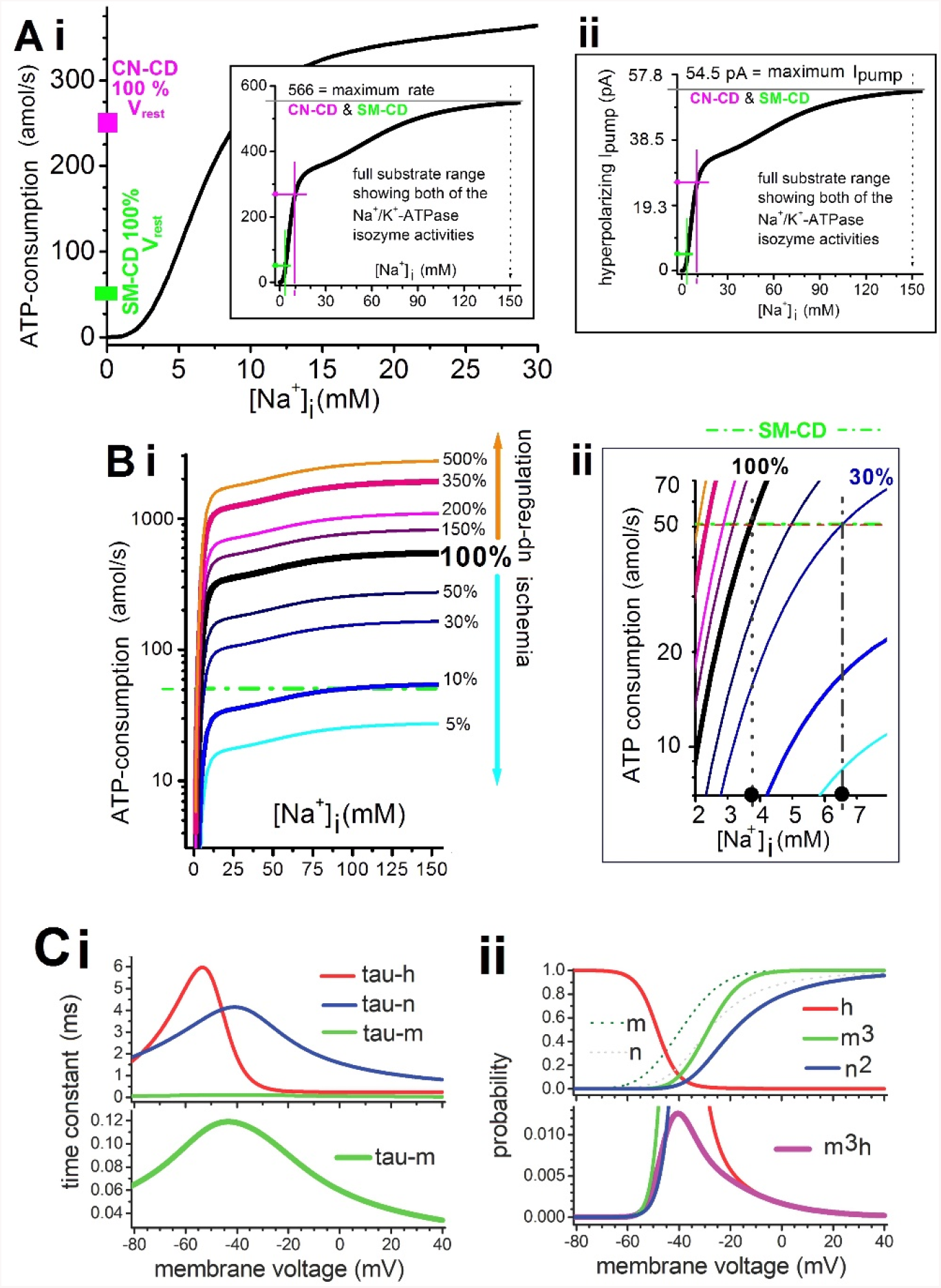
Pump and Excitability. **A,B. The pump: ATP-consumption (or hyperpolarizing I_pump_) as [Na^+^]_i_ varies** **Ai**. For the pump (Hamada et al 2003) used in all models here, ATP-consumption as a function of [Na^+^]_i_; normal pump-strength for CN-CD and SM-CD as indicated on two scales (0-30 mM, and inset, 0-160 mM). **Aii** replots the inset, but for hyperpolarizing I_pump_, showing its direct proportionality to ATP-consumption. **Bi**. Using a logarithmic Y-axis, ischemia and pump up-regulation are depicted by scaling the 100% pump-strength curve as labeled. **Bii**. Lower left region **Bi**, expanded. Typically for enzyme kinetics plots, the abscissa is the independent variable (“Given [Na^+^]_i_, what is the ATP-consumption?”), but for P-L/D systems at steady-state, the pump kinetics plot used “in reverse” gives a read-out of steady-state [Na^+^]_I_ as illustrated here for SM-CD and LA-CD. In the P-L/D systems, −I_Naleak_ =hyperpolarizing I_pump_ and so, via **Equation 19** a model’s I_Naleak_ converts to steady-state ATP-consumption. See dashed green line at 51.1amol/s (SM-CD) and almost-overlapping dashed red line at 50.7 amol/s (LA-CD); where these intersect their models’ pump-strength curve (100% for SM-CD; 30% for LA-CD), the X-co-ordinate (black dots, below) indicates steady-state [Na^+^]_i_. Likewise for CN-CD (X-axis crossing, magenta “crosshairs”, **Ai** inset). **Ci,ii. Nav and Kv channels’ electrical excitability in the P-L/D models** Hodgkin-Huxley (H-H) plots for the V-dependent channels used here, calculated using **Equation 7** and kinetic constants given in **Table 4**. The excitable (action potential competent) P-L/D models here provide explicit leak permeabilities (P_Cl_, P_K_ and P_Na_); there is, therefore, no non-specific “H-H gLeak”. Also unlike standard H-H formulations, driving forces are all electrodiffusive (not ohmic) (see **Methods, Equations 2** and **3**).

Generally-speaking, electrically-leakier systems deteriorate faster. Based on total leak permeabilities (see **Figure 8A**) SM*-CD (40.3 µm^3^/s) is electrically-leakier than CN*-CD (24.5 µm^3^/s). Why, then, the markedly slower anoxic rundown (gradient dissipation, V_m_ decay, swelling) of SM*-CD? Because in P-L/D systems, rate-limiting permeabilities govern rundown, not globlal electrical-conductivity (**Figure 3** legend provides more specifics). CN*-CD, with P_Cl_ small relative to (its major permeability) P_K_, represents “generic neuron” where, famously, the [small P_Cl_] slows anoxic swelling (Rungta et al 2015). Therefore, the physiologically noteworthy take-home from this rundown comparison is: in [big P_Cl_] SMFs, the extremely small P_Na_ confers substantially more safety from anoxic swelling than [small P_Cl_] does in [big I_Naleak_] neurons.

*A priori*, in P-L/D systems with unchanging permeabilities, a pump-strength sufficient to sustain steady-state gradients is also sufficient to restore systems after anoxic dissipation. **Figure 3** confirms this. SM*-CD, being less dissipated than CN*-CD at 105 minutes, recovers sooner, but whatever the rundown time, full recovery is assured (not shown).

### Electrophysiological lifestyles and excitable cell anoxia

Neurons can respond sensitively to small incoming currents because their input impedance is relatively large and their V_rest_ lies close to firing threshold. SMFs, by contrast, keep V_rest_ hyperpolarized far below firing threshold and their smaller input impedance (from [big P_Cl_]) prevents firing except in response to large excitatory postsynaptic currents (EPSCs) that, in situ, occur infrequently.

During anoxia (**Figure 4**) the behaviors of the P-L/D systems adapted for these distinctive “electrophysiological lifestyles” differ to an extreme degree; neurons perish rapidly and SMFs remain safe for prolonged periods. **Figure 4** follows CN-CD and SM-CD anoxic rundowns for 50 min (details in legend). Post-anoxic restoration is addressed elsewhere (for CN-CD, see Dijkstra et al (2016) and for SM-CD, see Morris et al submitted); suffice it to say that in both models post-anoxic recovery occurs only if I_pump_ is restored before ectopic firing ceases.

Instantaneous pump-off depolarization in CN-CD depolarizes V_m_ to firing threshold (**Figure 4A**). As ectopic APs fire for ∼1 minute, Nav and Kv channel opening precipitously dissipates the cation gradients. In CN-CD, a pathological voltage-activated Cl^-^ channel (Rungta et al 2015) hastens swelling. At at 50 minutes, CN-CD is 3X its initial volume (versus 2X for CN*-CD).

50 minutes into anoxia, the condition of SM-CD (**Figure 4B**) is equally disastrous. Having generated a half minute ectopic firing burst, it too has swollen 3X (versus only 1.2X for SM*-CD). The massive cation fluxes via Nav and Kv plus concurrently a catastrophically large [Cl^-^+H_2_O] influx. A different story is evident, however, when comparing systems, say, 20 minutes into anoxia. For 22.7 minutes, anoxic SM-CD avoids the Donnan-mediated swelling catastrophe, its prolonged rundown thanks to its hyperpolarized V_rest_ plus [small I_Naleak_]. Ancillary to [small I_Naleak_], moreover, its pump-off depolarization is small.

SMFs require [big P_Cl_] as part of their no-added-cost “low-internal-resistance Donnan battery”; the feature keeping V_m_ near V_rest_ during (non-EPSC) current perturbations. Until firing threshold is reached, that [big P_Cl_] is unproblematic. During those 22.7 minutes, in spite of [big P_Cl_], the minimal P-L/D system keeps Cl^-^ influx small because Cl^-^ feels no driving force at t=0 (steady-state E_Cl_=V_rest_ and during rundown E_Cl_ tracks V_m_). Though SM-CD’s P_K_ is small (compared to CN-CD), P_K_>>P_Na_, so Na^+^ entry through P_Na_ is mostly neutralized by K^+^ exit. With Cl^-^+H_2_O entry minimal, SM-CD at 22.7 minutes into anoxia has swollen a mere 1.05X. The tightly coupled Cl^-^+H_2_O entry over this period is plotted at lower left, **Figure 3B**.

In SM-CD, anoxic depolarization from −75 mV→ −65 mV takes 12 minutes. This accords well with the report (Clausen and Flatman 1977) that in ouabain-poisoned rat soleus fibers V_m_ runs down at ∼10 mV/10 minutes.

In SM-CD, a ±10-fold ΔP_Cl_ does not affect V_rest_ and rundown at 10X P_Cl_ and 0.1X P_Cl_ takes 23.8 and 20.7 minutes to ectopic firing, respectively. Thus, its importance in stabilizing SMF V_rest_ notwithstanding, [big P_Cl_] does not influence the value of V_rest_ (see **Figure 10**) and has little impact on its decay. As further evidence of the primacy, here, of [small I_Naleak_]: a mere 27% P_Na_ decrease (→V_rest_=-90 mV), as illustrated in Morris et al submitted, lengthens rundown-to-ectopic firing from 22.7 to 34 minutes.

**FIGURE 10.**
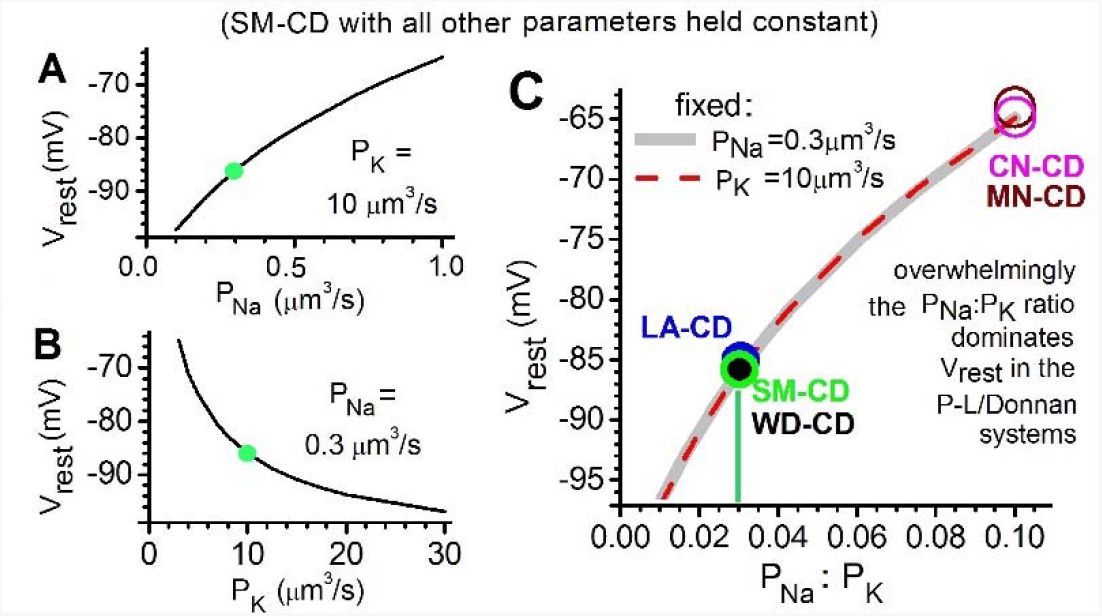
Cation Permeabilities and V_rest_. With SM-CD’s other parameters held constant, **A** and **B** plot V_rest_ as P_Na_ or P_K_ vary (green dots, normal SM-CD V_rest_) as they might, say, as a consequence of hormonal modulation. In **C**, plots **A** and **B** coalesce, giving V_rest_ as P_Na_:P_K_ varies. Green cross-hairs (P_Na_:P_K_ = 0.3) indicate SM-CD normal V_rest_ (−86.0 mV) with other P-L/D models V_rest_ also indicated at the P_Na_:P_K_ appropriate for each model. The models vary enormously in their PCl values, and CN-CD (but not MN-CD) has a K/Cl cotransporter that modifies E_Cl._ Nevertheless, V_rest_ is overwhelmingly dominated by the P_Na_:P_K_ ratio. These calculations all assume normal pump strength. As per **Figure 7Ai**, increasing SM-CD pump-strength above the normal (100%) negligibly affects V_rest_ but if pump strength falls below ∼20% V_rest_ depolarizes (due to diminished hyperpolarizing I_pump_ plus depolarizing Nav window current).

### WD-CD: a counterfactual Pump-Leak dominated SM-CD analog

Many key physiological traits of SM-CD depend directly on [small I_Naleak_] independent of [big P_Cl_]: SM-CD’s low resting ATP-consumption and consequently its large pump-reserve (ratio of maximal to resting ATP-consumption), its strongly hyperpolarized V_rest_, its slow anoxic rundown. It can be easy to lose sight of what [big P_Cl_] contributes to Donnan dominated ([big P_Cl_][small I_Naleak_]) ion homeostasis.

To illustrate quantitatively, we use WD-CD (Weak Donnan, **Tables 1** and **2**), a counterfactual Pump-Leak dominated analog of SM-CD with the same input impedance and V_rest_ (−86 mV). Its assigned P_Cl_ is that of a physiologically-bonafide Pump-Leak dominated system (i.e., CN-CD) and its P_Na_:P_K_ ratio is that of SM-CD, but with larger absolute values to attain the input impedance of SM-CD. In counterfactual WD-CD, therefore, P_K_ (not P_Cl_) is the major contributor to low input impedance. Like SM-CD, WD-CD is a minimal P-L/D system. Its [big I_Naleak_] puts resting ATP-consumption at 3.5X that of SM-CD; its pump-reserve is only 3.1-fold. Though spending 3.5X more for steady-state Na^+^ extrusion than SM-CD, its [Na^+^]_i_ is almost twice that of SM-CD, leaving it less able than SM-CD to support SMF’s E_Na_-dissipating Na^+^-transporters.

In [big P_Cl_] SM-CD, excitability modulation is straightforward (ΔP_Cl_ leaves ATP-consumption and V_rest_ unchanged) but WD-CD modulation is complex. Altering its input impedance via ΔP_K_→ΔV_rest_. Modulating it with no ΔV_rest_ requires changing P_K_ and P_Na_ at fixed ratio. Any ↑P_Na_→↑ATP-consumption.

Ischemic episodes occur in the normal course of events for SMFs, making SM-CD’s exceptionally slow anoxic rundown a “time-is-money” benefit. Slow rundown makes it likelier that blood-flow resumes before ectopic firing triggers damaging contractures and cytotoxic swelling. **Figure 5** shows WD-CD anoxic rundown. Thanks to [small P_Cl_] and initially zero Cl^-^ driving force, swelling prior to ectopic-APs is inconsequential (a neuron-like attribute). Nevertheless, due to its [big I_Naleak_] WD-CD rundown-to-firing is 4X faster than in SM-CD (−75→-65 mV in 2.9 minutes versus 12 min for SM-CD). Exacerbating this perilous rapidity is WD-CD’s 5.5 mV pump-off depolarization (versus <2 mV in SM-CD).

### Evolution chose Donnan dominated ion homeostasis for SMFs

Like a neuron, a “WD-CD SMF” must continually “pay the pump” a) for its costlier V_rest_ and b) for most (i.e., the non-P_Cl_ portion) of its resting conductance. Like a neuron, WD-CD, is vulnerable to ischemia. An organism using Pump-Leak dominated “WD-CD muscles” for all its mostly-quiescent SMFs would chronically elevate its whole-body expenditure, jeopardizing fuel supplies for contractility. **Figure Sup01C** shows that WD-CD (see the ATP-consumption trace) handles the 1200 AP stress-test as well as SM-CD. Its base-line energy consumption, however, is >3.5X higher.

For neurons and the whole-organism that supplies their ATP, payback for use of a costly Pump-Leak dominated system is their agile, nuanced integration of diverse information-rich electrical inputs. This merits the expenditure and the added vulnerability. By leaving SMF decision-making (when to contract?) entirely to remotely-located groupings of high-cost neurons, vertebrate SMFs could benefit from their robust low-cost Donnan dominated homeostasis strategy, allowing more ATP for contractility.

SMFs’ [small I_Naleak_] checks treacherous Donnan forces, while fostering the maintenance of a large pump-reserve and a strongly hyperpolarized, exceptionally low-cost V_rest_ that, thanks to SMF’s “pre-paid” Cl^-^-battery (i.e. [big P_Cl_]+[Donnan effectors]), is not readily perturbed.

### The slow ion homeostatic response to a fast SMF electrical event

The canonical ionic perturbation of SMF ion homeostasis is a cholinergic motorneuron-triggered AP (**Figure 2B**). For 3 minimal P-L/D models, Donnan dominated SM-CD (green), its ischemic (30% pump-strength) variant, LA-CD (blue), and (counterfactual) Pump-Leak dominated WD-CD (red) (see **Tables 1** and **2**), **Figure 6** details the response to an EPSC-triggered AP. Here, overlying the SM-CD AP of **Figure 2B** are almost-identical LA-CD and WD-CD APs. Time-axes and the various Y-axes are varied to resolve fast AP-related perturbations and the slower P-L/D responses. Single-AP ion and volume responses would be immeasurably small, but **Figure 2A,B** showed (for SM-CD) that the cumulative perturbation from 1200 APs elicits larger-magnitude P-L/D responses of nearly the same duration as for one AP. By 1200-APs in SM-CD (**Figure 2A**), [Na^+^]_I_ rises almost 25 mM above steady-state whereas post-one-AP, [Na^+^]_I_ increases <0.05 mM. For contraction-inhibited SMFs (current or voltage-clamped), a new 4-microelectrode technique enables simultaneous monitoring of multiple [ion]_i_ (Heiny et al 2019; DiFranco et al 2019; for measurably-large perturbations, an AP train (not a single AP) would be needed).

**Figure 6** dissects the underlying P-L/D processes.

The ATP-consumption trajectories are straightforward: after rapid double-jumps (responding to ↑[Na^+^]_I_ from the EPSC and AP), pump activity decays monotonically to steady-state over several minutes. WD-CD’s trajectory is off-scale near 180 amol/s but otherwise like that of SM-CD (see Y-axis ranges, double-jump boxes) and so the Na^+^-extrusion and K^+^-uptake (difference) trajectories for SM-CD and WD-CD overlap, while weaker pump-strength LA-CD lags.

Return to steady-state [Cl^-^] and Vol_cell_, rather than being monotonic, involves a single oscillation, reflecting the interplay of P-L/D systems’ two sensor/effector processes. Once AP channels close, V_m_(t) reflects hyperpolarizing I_pump_(t). After 1200 APs in SM-CD (**Figure 2A;** see also **Figure Supp01B,Cii**), V_m_(t) hyperpolarizes and temporally aligned oscillations of [Cl^-^]_i_(t) and Vol_cell_(t) occur as in **Figure 6**, but writ large.

All 3 models oscillate, their distinctive features revealing the forces at work (detailed in legend). Why the oscillation? Open AChR channels support mostly Na^+^ entry, then Nav channels open before Kv channels. Immediately after any excess Na^+^ entry (see AP-induced H_2_O blip), cells are Na^+^ and H_2_O and Cl^—^overloaded. [Na^+^]_i_-stimulated electrogenic I_pump_ extrudes 3Na^+^ per 2K^+^ imported, slightly hyperpolarizing the V_m_ (top panel, right). Shortly, the ongoing Na^+^-extrusion makes the swollen cells shrink below normal because the pump keeps expelling 3Na^+^ per 2K^+^ imported, and neutralizing-Cl^-^ plus osmo-balancing-H_2_O accompany the extruded Na^+^ (i.e. they exit the cell). [Cl^-^]_i_ and Vol_cell_ thus drop below their steady-state levels until, as [Na^+^]_I_ continues falling, I_pump_ and I_Naleak_ equalize and steady-state is re-attained.

The ion homeostatic feedback oscillation of Vol_cell_(t) and [Cl^-^]_i_(t), we call a “Donnan bounce”. The Vol_cell_(t) panel (right) shows that the Donnan bounce amplitude of Pump-leak dominated WD-CD is <1/10^th^ that of Donnan dominated SM-CD. A P-L/D system’s Donnan bounce is bigger when P_Cl_ is bigger and slower when pump-strength is weaker.

Based on the **Figure 6** single-AP responses, one might ask:

**1**) Since WD-CD performs well post-AP, why would Pump-Leak dominated neurons (say) not go a step further, eliminate Cl^-^ channels, and use Pump-Leak homeostasis? The answer: this could work only if AP-related Na^+^ and K^+^ fluxes and the Na^+^ and K^+^ driving forces were precisely adjusted to yield zero excess Na^+^ entry (AP-induced “H_2_O blips” would drop fully) AND if pumps were electroneutral (electrogenic pumping polarizes V_m_, altering any perfectly-tuned driving forces). This zero-P_Cl_ system would have zero physiological latitude.

**2**) WD-CD experiences a smaller H_2_O blip and thus a smaller Donnan bounce; is that not preferable? Inside a cranium, ΔVol_cell_ is undesirable, but for SMFs the ΔVol_cell_ of SM-CD is vanishingly small, physiologically-speaking, whereas WD-CD’s chronic 3.5X↑ATP-consumption is an untenably large differential that, for low-duty-cycle SMFs, disqualifies the Pump-Leak dominated analog even ignoring its overly-fast anoxic rundown (**Figure 5B**).

### SM-CD, ion homeostatic thresholds and bistability

SMF pump-strength varies up/down through hormonal modulation and misadventure (interrupted blood flow, toxins, sarcolemma bleb-damage) (Pirkmajer and Chibalin 2016). Given SMF’s interacting non-linear Na^+^-flux mechanisms (I_pump_([Na^+^]_i_) and I_Nav_(V_m_) (**Figure 9A,C**), and the non-linear driving forces acting on permeant ions, it is to be expected that fibers’ P-L/D steady-states would vary non-linearly with pump-strength. Bifurcation plots for SM-CD (**Figure 7**) encapsulate these non-linearites.

For neurons in general and CN-CD in particular (Dijkstra et al 2016), it was shown (Hubel et al 2014) that pump-strength changes cause smoothly-varying (albeit non-linear) changes in steady-state values, until a system-specific unstable threshold is encountered. Here we examine this property for SM-CD. **Figure 7Aii** is the SM-CD destabilization trajectory (normal-pump-strength→saddle-node-pump-strength). Once spontaneous destabilization (ectopic firing) commences, the minimal-pump-strength system quickly degrades then slowly continues (not to DE, since pump-strength>0) to a “DE-like” pathological steady-state. SM-CD can now only return to a physiological steady-state (**Figure Sup02 Ci,ii,iii**) if it up-regulates its pump-strength (=↑↑hyperpolarizing I_maxpump_) enough to repolarize itself to a different unstable threshold (→more ectopic firing) from which (post-APs) it repolarizes to a physiological steady-state. This bistable two-thresholds scenario (“ion homeostatic excitability”) is a feature of excitable P-L/D systems (Hubel et al 2014; Morris 2018).

The parameter-specific bifurcation plots for a P-L/D system coincide; to know SM-CD’s steady-state V_m_, [Na^+^]_i_, Vol_cell_, etc at, say, 10% pump-strength, consult each plot at that X-axis value. Furthermore, SM-CD/CN-CD comparisons are informative because their 100% pump-strength and C_m_ values are identical. For SM-CD, the lowest pump-strength able to sustain a physiological steady-state is marginally <8% (the saddle-node is ∼7.9%). Comparison of saddle-node pump-strengths (∼8%(SM-CD), ∼65%(CN-CD)) re-emphasizes the impressive ion homeostatic robustness of SM-CD. Due to 3X↑Nav density, V_**m**_ at SM-CD’s saddle-node is less depolarized than for CN-CD (**Figure 7Ai**); but note that SM-CD, at its saddle-node, though profoundly Na^+^-loaded, is hardly swollen (**7Bi,Cii**). Thus, unlike CN-CD, SM-CD can be profoundly ischemic and Na^+^-loaded with very little swelling. In humans, various methodologies including ^23^Na-proton-MRI, point to “non-osmotic Na^+^-loading” in connection with myotonia (Weber et al 2006), over-exertion pain (Yu et al 2013), anaerobic exertion (Hammon et al 2015), and the chronically Na^+^-overloaded fibers of DMD patients (Gerhalter et al 2019).

The Vol_cell_ plot (**Figure 7Ci**) emphasizes the far-from-equilibrium state of healthy cells and the potency of the Donnan forces held in check by ion homeostasis by plotting the full range between DE and physiological Vol_cell_ (black line) levels. **7Cii,iii** then bring the focus to the physiological region. Very little swelling is tolerated by CN-CD, but even SM-CD, whose saddle-node is reached at ∼2088 µm^3^, swells only ∼5% (↑Vol_cell_). Working SMFs can safely withstand greater substantial swelling at healthier pump-strengths with Na^+^-transporters operative (Usher-smith et al 2009; Lindinger et al 2011).

The **Figure 7Aii** trajectory follows SM-CD to a degraded ATP-consuming pathological steady-state. If (implausibly) a SMF survived the ectopic firing/contractures and stabilized there, it would occupy a pathological steady-state point on the continuum (blue) directly above the black **X**. Recovery would require this degraded system to up-regulate its pump-strength to **#**. For completeness, **Figure Sup02Cii,iii** shows SM-CD’s V_m_(t) trajectory when pump-strength is increased from 338% to 339%(#), whence it destabilizes and converges on the physiological steady-state continuum (directly below **#**).

Though peril clearly awaits if a saddle-node is encountered, ischemic SMFs are profoundly safer than ischemic neurons. If CN-CD drops 100%→65%, only doubling its pump-strength (pink #) will save it. By contrast, SM-CD’s physiological continuum extends down to 8%; over this entire ischemic range, any pump-strength improvement, no matter how small, moves the system smoothly back towards 100% (**Figure 7Bii**). SMFs commonly experience transient benign ↓blood-flow (e.g., from sitting on one’s foot). The slowness of anoxic/ischemic rundown times (**7Aii** inset), together with **7Bii** (recoverability) signifies that resting SMFs are well adapted to prolonged deep ischemic bouts (provided bouts end before myotonic firing commences and/or before motorneurons trigger APs).

Bifurcation analysis of SM-CD is congruent with the clinical situation, compartment syndrome, where, a normally-robust system harbours a treacherous threshold. Compartment syndrome describes progressive potentially life-threatening (gangrene) muscle ischemia, usually involving a limb, typically post-trauma. Johnstone and Ball (2019) indicate that “membrane injury and subsequent significant dysfunction is the central factor leading to irreversible {SMF} injury” and emphasize that because SMFs’ remarkable tolerance of progressive ischemia ends abruptly, a fundamental requirement for better diagnoses is a means (intramuscular pH, perhaps) of identifying the ischemic threshold.

### SM-CD dynamic range: up-regulated pumps and/or down-regulated P_Na_

Intermittently, locomotory, thoracic, vocal, diaphragm and other muscles experience upsurges of ion homeostatic demand. **Figure 2A** showed a realistic hyper-stimulation of V-gated channels. *In situ*, Na^+^-transporters too would need to be supported. As Fraser and Huang (2004) show, use of NKCC, a cation-chloride cotransporter, would ↑[Na^+^]_I_, ↑[Cl^-^]_i_, ↑Vol_cell,_ ↓[K^+^]_i,_ depolarize V_m_ and ↑I_pump_. To extend its physiological dynamic range for excitability and transport, SM-CD shows us, a SMF could optimize its ATP consumption as described next.

Embedded in the physiological steady-state [Na^+^]_i_ continuum, **Figure 7Bi**, is an easily-overlooked P-L/D feature germane to physiological dynamic range. That plot’s “boring” flattish region implies that no metabolic penalty is incurred for keeping pump-density chronically up-regulated (i.e., extra-large pump-reserves). Large ↑pump-strength has little affect on [Na^+^]_i_ (**Figure 7Bii**). Increasing SM-CD pump-strength 100→200% (pump-reserve=11→22-fold, a level reported for rat soleus (Clausen 2015)) decreases [Na^+^]_i_ to 2.9 mM (**7Bii**), hyperpolarizes V_rest_ minimally (**7Ai**) leaving ATP-consumption at 51.1 amol/s (**Figure 9Bii;** X=2.9 mM, 200%-curve Y-axis intersect). **Figure Sup01A** illustrates that pump-reserve=22-fold accelerates post-stress-test recovery.

Temperate-climate amphibians and reptiles confront “scheduled” (i.e., seasonal) bouts of extreme ischemia. This requires modulation of excitable cell ion homeostasis (Pérez-Pinzón et al 1992; Donohue et al 2000). A testable prediction emerging from SM-CD is that, prior to entering hibernation, hibernators would increase SMF pump-density. Following the deep super-prolonged ischemic rundown of hibernation, this would expedite springtime recovery. If borne out, regulatory signaling systems could be sought. Another prediction: hibernators would simultaneously reduce P_Na_ for reasons outlined next.

A ↓P_Na_ would improve a SMF’s dynamic range. Why? Because ↓P_Na→_↓I_pump_. It also counteracts Na^+^-cotransporter-induced depolarization. For example, reducing P_Na_ from 0.3 µm^3^/s to 0.22 µm^3^/s hyperpolarizes SM-CD V_rest_ by 4 mV (to −90 mV; see **Figure 10**). Because ↓[small I_Naleak_]→↓I_pump,_ ATP-consumption drops ∼25% (51.1 amol/s→38.7 amol/s). This hypothetical change illustrates, yet again, the energetically pivotal role of SMFs’ [small I_Naleak_] and hence of their small P_Na_. Had we picked −90 mV as V_rest_ for SM-CD, the SM-CD/CN-CD ATP-consumption comparison would have been 1/7.0 instead of 1/5.3.

### SMFs’ small P_Na_ in GHK versus P-L/D settings

The immense physiological importance of SMFs’ small P_Na_ has gone largely unrecognized. It is understood that V_rest_ depends on P_Na_:P_K_. It is understood that P_Na_ is the conduit for I_Naleak_, the determinant of I_pump_(=-I_Naleak_). It is understood that P_Na_ thereby establishes the pump’s small direct contribution (several millivolts) to V_rest_ (e.g., Sperelakis 2012). However, in spite of Fraser and Huang (2004)’s thorough exposition, it is under-recognized that for cells performing ion homeostasis (living cells) the zero-current Goldman-Hodgkin-Katz (GHK) equation is an inappropriate V_rest_ descriptor. This is not biophysical nit-picking.

Decreasing a small P_Na_ in a P_K_>P_Na_ GHK setting simply renders it increasingly inconsequential, and P_K_ more dominant. By contrast, in a P-L/D setting, as just illustrated for SM-CD, decreasing an already-small P_Na_ has physiologically consequential ramifications: ↓P_Na_→↓ATP-consumption and ↑pump-reserve. And in a transiently anoxic P-L/D system, ↓P_Na_→slower rundown and then faster recovery.

### The unidentified P_Na_ of SMFs

In SMFs, P_Na_ (and ΔP_Na_) has impressive physiological heft, but what is P_Na_? Neurons, smooth muscle and pancreatic cells use Na-Leak-Channel-Nonselective (NALCN) as a hormonally-regulated P_Na_ (Lu et al 2007; Senatore and Spafford 2013; Cochet-Bissuel et al 2014; Lutas et al 2018; Philippart et al 2018), but to date, NALCN has not been found in SMFs. Identifying the SMF P_Na_ and understanding its variability is considered critical in connection with myotonia (Metzger 2020). Persistent Nav channels (Gage et al 1989) might contribute, but since tetrodotoxin does not hyperpolarize V_rest_ in healthy fibers (e.g., Pickar et al 1991) this seems unlikely. The SMF P_Na_ could, like NALCN, be a cation channel (see Morris et al submitted). SMFs have several identified cation channels (see Metzger et al 2020) plus their routinely-detected unidentified mechanosensitive cation channel (Guharay and Sachs 1984; 1985; not a Piezo-channel (Suchyna 2017)). A precedent is worth recalling here: *Aplysia* neurons’ serotonin and arachidonic acid regulated resting K^+^-channels (Belardetti et al 1986) are adventitiously mechanosensitive (e.g., see Vandorpe et al 1994; Morris and Horn 1991; Methfessel et al 1986; Zhang and Hamill, 2000; Zhang et al 2000; Morris 2012). Likewise for SMF mechanosensitive cation channels, but decades on, they have no attributed function (Lansman 2015; Suchyna 2017). A hypothesis worth pursuing: this cation channel does for SMFs what NALCN does for smooth muscle (e.g. Reinl et al 2018). Jointly, P_K_ and P_Na_ establish V_rest._ Another critical and unresolved issue for SMF ion homeostasis, therefore, is whether/how they are jointly modulated (Kuba and Nohmi 1987; Donohue et al 2000; DiFranco et al 2015). Molecular identification of P_Na_ in SMFs could empower broad-reaching lines of biological and biomedical inquiry.

## DISCUSSION

### Ion homeostasis needs Donnan effectors

Comparative modeling of simple yet reassuringly realistic P-L/D systems here shows that, to serve distinctive electrophysiological lifestyles, SMFs and neurons evolved distinctive ion homeostatic strategies. Neuronal ion homeostasis biophysics was recently explained (Dijkstra et al 2016), but notwithstanding the foundational Fraser and Huang (2004) paper, SMF ion homeostasis mostly gets by-passed. For animal cells in general, moreover, the physiological necessity of [Donnan effectors + P_Cl_] for achieving ion homeostasis remains under-appreciated (Dmitriev et al 2019; Kay 2018).

To clarify the integral role of Donnan effectors in ion homeostasis, we outline first principles -- set point, sensor/effector feedbacks (**Figure 1A**)-- in their evolutionary context. This reveals ion homeostasis as a Pump-Leak/Donnan process. We depict “minimal” P-L/D systems at steady-state (**Figure 1B**), characterizing those P-L/D systems permeable mostly to anions as Donnan dominated, and those permeable mostly to cations as Pump-Leak dominated.

Computationally meshing the Fraser and Huang (2004) SMF model and the more accessible (compare pump descriptors) model of cortical neurons (Dijkstra et al 2017) we generated SM-CD, a minimal Donnan dominated excitable model for a SMF “ion homeostatic unit” (**Figure 8A**; **Tables 1, 2**).

Using SM-CD (and variant models) we demonstrate how low duty-cycle, low-excitability syncytial SMFs and more broadly, skeletal muscle, benefits from SMFs’ Donnan dominated strategy. The robust [big P_Cl_][small I_Naleak_] process makes an ion homeostatic/electrophysiologic virtue of the unavoidable “vice” of Donnan effectors. Neurons’ Pump-Leak dominated [small P_Cl_][big I_Naleak_] strategy, though far costlier and more imperiled by ischemia, is necessary for an information-processing lifestyle. For early vertebrates, the whole-organism contexts (**Table 2**) for these two strategies would have been an extreme version of the situation for humans: a low-cost Donnan dominated strategy for the body’s largest tissue and a costly pump-Leak dominated strategy for the small fractional mass of neurons. **Table 3** identifies and summarizes quantitatively several of the differences.

**Table 3.** Some operational attributes of the P-L/D models and new queries

|  | **Model name** | **SM-CD** | **WD-CD** | **CN-CD** |
| --- | --- | --- | --- | --- |
|  | *P-L/D model depicts* | “slice” of multinucleate SMF (estimated to be ~1 myonuclear domain volume) | a <u>counterfactual</u> “tool”; P-L dominated analog of SM-CD | a roundish soma of a cortical neuron in brain-slice (Dijkstra et al 2016) |
| 1 | *ATP-consumption (or, relative to SM-CD)* | 51.1 amol/s (1X) | 180.4 amol/s (3.5X) | 270.9 amol/s (5.3X) |
| 2 | *pump-reserve* | 11.1-fold | 3.1-fold | 2.1-fold |
| 3 | *rate-limiting P for $\Delta V(t)$ as anoxic rundown starts* | $P_{Na} (=0.3 \mu m^3/s)$ | $P_{Cl} (=2.5 \mu m^3/s, \text{ and, at } t=0, V_m = E_{Cl})$ | $P_{Cl} (=2.5 \mu m^3/s, \text{ and, at } t=0, V_m \text{ positive to } E_{Cl})$ |
| 4 | *time till ectopic firing during anoxic rundown* | 22.7 minutes (rundown at: 10 mV per 12 min) | 6 minutes (10 mV per 2.9 min) | <100 milliseconds (triggered by pump-off dep’n) |
| 5 | *no V-gated channels (\*) -- 20 min into anoxic rundown: $V_m, Vol_{cell}$* | (SM*-CD)<br>depolarized by 19 mV<br>swollen 1.05-fold | --- | (CN*-CD)<br>depolarized by 45 mV<br>swollen 1.4-fold |
| 6 | *saddle-node pump-strength & $[Na^+]_i$* | 7.9% pump-strength, $[Na^+]_i = 88 \text{ mM}$ | 26.5% pump-strength, $[Na^+]_i = 92 \text{ mM}$ | 65% pump-strength, $[Na^+]_i = 42$ |
| 7 | *patho-physio threshold pump-strength* | 339% pump-strength ($V_{pathological} = -34 \text{ mV}$) | ---- | 185% pump-strength ($V_{pathological} = -37 \text{ mV}$) |
| 8 | *examples of conditions to examine ion homeostatically* | hibernation onset/offset, compartment syndrome, delayed onset muscle soreness, Nav & other channelopathies, muscular dystrophies | (a WD-CD/SM hybrid) the “$P_{Cl}$ disease”, myotonia –mutant CLC-1 → excitable SMFs with [small $P_{Cl}$][small $I_{NaLeak}$] | ischemia, migraines, epilepsy (Dijkstra et al 2016), optimizing analgesic protocols for neuropathic pain (see Lorenzo et al 2020) |
| 9 | *examples of relevant physiological/biological/ evolutionary questions (re SM-CD and CN-CD)* | CLC-1 underlies [big $P_{Cl}$]. Does this channel’s very small unitary-g somehow benefit SMF ion homeostasis? | can smooth muscle (Lauf et al 2019) be modeled by modifying WD-CD ($Ca^{2+}$-gCl & gK, co-K/Cl)? | did early vertebrate hindbrain vascularization (Rahmat & Gilland 2014) prioritize fuelling of central neurons? |

### Ion homeostatic efficiency and SMF morphology

Even beyond **Table 3’s list of** CN-CD/SM-CD comparisons, ancestral vertebrates would have benefitted enormously from Donnan dominated ion homeostasis. This assertion assumes that then, as now, SMFs were syncytial (mammal SMFs have hundreds to thousands of nuclei (Ganassi et al 2018; for myonuclear domains, see Ross et al 2018). SM-CD and CN-CD have the same 2000 µm^2^ surface area (SA) (C_m_) and volume (**Figure 8A,Bii,Cii**). In situ however, SM-CD would not be mononucleate. It would be one of many cross-sectional slices of a “cylindrical” SMF comprised of hundreds to thousands of such slices (**Figure 8C,D**). By spherical geometry, 2000 µm^2^ can fully enclose at most 8419 µm^3^, while by cylindrical geometry, the maximal volume encircled by 2000 µm^2^ varies inversely with “ring”-width; for example, disposed as a 10 µm slice, 2000 µm^2^ of sarcolemma (no slack) encircles ∼32,000 µm^3^ (see **Figure 8Di,ii**).

Syncytial morphology thus increases the cytoplasmic volume served (ion homeostatically) by a “unit” of sarcolemmal SA. SMFs are usually at steady-state, and crucially, steady-state ATP-consumption is independent of SA/Vol_cell_. SA/Vol_cell_ does, however, matter during ionic perturbations: scaling linearly, for fixed-SA, both anoxic rundown times and ion homeostatic recovery times lengthen with ↑Vol_cell_. For low duty-cycle SMFs, the advantages conferred by slower rundown would, overall, substantially outweigh any drawbacks of slower recovery.

### The SM-CD “ion homeostatic unit” and myonuclear domain volume

For mouse, data from Harris et al (2005); Mantilla et al (2008); Egner et al (2016) indicate: SMF radius ∼15-30 µm, myonuclear density ∼50/mm, myonuclear domain volume 15,000-30,000 µm^3^. For mouse SMFs, SM-CD’s 2000 µm^2^ “ion homeostatic unit” would encircle more myoplasmic volume than depicted in **Figure 8Bii**. If mouse SMF nuclear density is ∼1 nucleus/20 µm (∼50 nuclei/mm→∼0.5 nucleus/10 µm), a plausible “volume-corrected” SM-CD would be a 20 µm slice of 15 µm radius. If it had SA=1885 µm^2^ plus a slack 115 µm^2^ for passive tension-buffering (1.06-fold swelling) that would be 2000 µm^2^ encircling 14,137 µm^3^. (Sarcolemma’s dynamic tension-buffering (Sinha et al 2011; Lo et al 2016; Morris 2018) is not modeled here). A SM-CD unit such as this would be serving ∼1 myonuclear domain.

The 2000 µm^2^ “ion homeostatic unit” of SM-CD was picked with no consideration of myonuclear domain volume, but in retrospect, it is reasonable that a roughly comparable unit of SA would serve (and be serviced by) one nucleus in both neurons and SMFs. For neurons, one nucleus serves soma, dendrites and axon, except when neurons become “giants” and either massively amplify their DNA-content in one nucleus (Lasek and Dower 1971) or become multinucleate (Mackie 1990).

Thus for these radically different cytomorphologies, the SA value taken from neurons to facilitate inter-model ion-flux comparisons seems roughly appropriate to associate with one myonucleus, but this remains a speculation; myonuclear domain volumes are routinely reported (e.g. Schwartz et al 2016) for SMFs, but we found no data on sarcolemmal-SA per myonuclear domain.

### Switching off excitability for SM-CD robustness

An elevated risk accompanies the energetic benefits of syncytial morphology: loss of thousands of nuclei from one sarcolemmal tear. SMFs mitigate this danger via impressively reliable repair mechanisms (Andrews et al 2014; Barthélémy et al 2018; Manoharan et al 2019). Almost certainly the large pump-reserve facilitated by SMFs’ Donnan dominated strategy would be critical for re-establishing ion gradients post-tear. Imperative too would be diminished excitability. Whereas inexcitable SM*-CD recovers fully after >250 minutes of anoxic-rundown, at 25 minutes, excitable SM-CD having fired ectopically has swollen catastrophically. In the contexts of both ischemia and sarcolemmal tearing (McElhanon and Bhattacharya 2018; Hotfiel 2018), entry of Nav1.4 channels into slow-inactivated states (Webb et al 2009) needs to be studied.

### P-L/D steady-states for real cells

“Set point” for ion homeostatic systems is a particular collection of steady-state values {Vol_cell_, V_m,_ [ion]_i_} (**Figure 1A)**. “Set point” may sound like a system “goal”, but we reiterate, deviations from “set point” values are not what is sensed operationally in a perturbed P-L/D system; the {Vol_cell_, V_m,_ [ion]_i_} value-set is the CONSEQUENCE, not “the goal” of the P-L/D system’s feedback processes.

Clarity here makes it easier to see how, in large real cells (unlike in models) non-negligible distances and real diffusion times will yield “fuzzy” steady-state value-sets without abrogating the P-L/D concepts. Consider a quiescent 5 cm sartorius SMF or a 50 cm motorneuron. There is ad libitum ATP, axial resistance, and the densities of leak-channels and pumps are inhomogenous. All along the cell, the P-L/D systems’ two sensor/effector mechanisms (**Figure 1A**) are continually sensing/effecting. According to local densities of P_Cl_, P_Na_, P_K,_ pump, [Na^+^]_I-_sensing by the pumps affects I_pump_ and simultaneously, deviations from osmo-balance and compartment neutrality are stochastically/electrostatically sensed/countered via passive H_2_O and Cl^-^ fluxes. At, say, the axon terminal, “V_rest_” will likely differ from “V_rest_” 50 cm away at the soma. Thus, at steady-state there would be standing [ion]_I_ gradients and steady-state axial currents. In quiescent SMFs, with their endplate specializations and extensive t-tubular invaginations contiguous with the sarcolemma (Pedersen at al 2001; Fraser et al 2011), true steady-state will likewise be spatially inhomogeneous. Rhythmically-active excitable cells’ ion homeostatic steady-states are limit cycles (see Cha and Noma, 2012); thus, P-L/D set points can also be inhomogeneous in the temporal domain.

### Excitable cells and P-L/D biotechnology

Symmorphosis (Fogarty and Sieck 2019), the evolutionary concept of economy of biological design, predicts that structural properties will be matched to functional demands. To analyze excitable cell ion homeostasis via a comparable conceptual filter, an ideal case would be the electric fish *Electrophorus electricus* (Gallant et al 2014; Catania 2015, 2019), given its low-duty-cycle syncytial SMFs and (derived evolutionarily from them) 3 subclasses of high-duty-cycle syncytial electrocytes (Traeger et al 2017). Available molecular (e.g., Nav channels, Kv, pumps) and morphological data for both cell types (Ching et al 2015, 2016; Thornhill et al 2003; Schwartz et al 1975; Marchado et al 1983) could be augmented with comparative proteomics (e.g., how much if any ClC-1 is expressed in electrocytes?). Molecular evolution data are available for O_2_-delivery (globin-dependent) to SMFs and the electrocyte sub-classes (Tian et al 2017). Questions of ion homeostatic design economy/robustness/agility are intriguing from the perspective of evolutionary biology, and also increasingly, in the context of creative new biomimetic ventures such as the bioinspired engineering of “soft power sources” (Schroeder et al 2017) and the development of optogenetic skeletal muscle bioactuators and resilient self-healing muscle-machine interfaces (Raman et al 2016, 2017, 2019).

### Prospects

Recently-developed cell-physiological techniques (Heiny et al 2019; DiFranco et al 2019) could make it possible to obtain concurrent SMF [ion]_i_ data (plus V_m_(t)) along the lines of **Figure 2A**, while for whole tissue in humans, MRI advances are allowing for the non-invasive probing of ion homeostatic parameters in healthy, injured and diseased subjects (e.g., Hammon et al 2015; Dahlmann et al 2016, Gerhalter et al 2017, 2019; Zhang et al 2020). The accessible theoretical framework process presented here for SMF ion homeostasis as a P-L/D process should help guide investigative directions and facilitate interpretation of new data streams.

Though SM-CD is radically simple, adding and modifying components as appropriate is entirely feasible. A little-exploited but thoroughly rigorous cardiomyocyte CD-approach model (Cha and Noma 2012) demonstrates that even with a plethora of channels, transporters, compartments and binding reactions, a properly-designed CD-approach model will converge on steady-state. But, even in its present minimal state, SM-CD has provided comparative physiological insight relevant to vertebrate evolution. It also helps explain how the ischemic membrane-damaged SMFs of DMD patients can survive for decades (Morris et al submitted). Given appropriate refinements, SM-CD modeling should be helpful in connection with muscle fatigue, endurance, trauma, volume regulation, ischemia-reperfusion injury, ion channelopathies, hibernation, cold tolerance, sarcopenia and more (e.g., Donohue et al 2000; Lindinger et al 2011; dePaoli et al 2013; Yu et al 2013; Clausen 2015; Ammar et al 2015; Bækgaard Nielsen et al 2017; Boërio et al 2018; Hostrup and Bangsbo 2017; Hotfiel et al 2018; Cannon 2018; Copithorne and Rice 2019; Li et al 2020; Metzger et al 2020; Altamura et al 2020; Surkar et al 2020; Thoma et al 2020).

## METHODS

### A Pump-Leak/Donnan (P-L/D) model for SMFs

Skeletal muscle fiber (SMF) ion homeostasis is modeled (SM-CD) by a single compartment limited by a semipermeable membrane in an “infinite” (fixed concentrations) extracellular volume (**Figure 8, Table 2**). SM-CD constitutes a single P-L/D “ion homeostatic unit”; an actual multinucleate SMF would be comprised of hundreds or even thousands of such units (depending on sarcolemma area; see **Figure Meth 01C,D**) would likely have a smaller SA/V (surface area to volume) ratio than SM-CD (thus, slower Δ[ion]_i_ dynamics). The SM-CD membrane encloses a fixed quantity of Donnan effectors: A^-^, impermeant monovalent anions. The extracellular medium has a small fixed A^-^ concentration. The membrane is permeable to Na^+^, K^+^, and Cl^-^, ions whose extracellular concentrations are fixed. SM-CD has the same physical characteristics as the Dijkstra et al (2016) neuronal cell, CN-CD (see **Table 2**): a resting cell volume (Vol_cell_) of 2000 µm^3^ and a constant C_m_ corresponding to a SA of 2000 µm^2^. As such, resting state cells are flaccid, i.e., not maximally inflated (**Figure 8**). Lipid bilayers tolerate little lateral expansion, but unless a model-cell inflated to spherical its membrane would not be subject to tension and rupture. The permeation pathways of SM-CD and CN-CD (given below) include “resting leak conductances” (i.e., permeation pathways) for 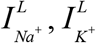, 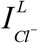, and 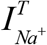 and voltage-gated conductances (permeation pathways) for a transient sodium current, 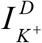, for delayed rectifier potassium current,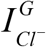, and (CN-CD only) a voltage-dependent chloride current, 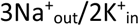. Driving forces acting on ions are, in all cases, electrodiffusive, as depicted by Goldman Hodgkin Katz (GHK) current equations (Hille, 2001). The same 3Na^+^_out_ /2K^+^_in_ ATPase pump model (Hamada et al 2003) is used throughout. It produces hyperpolarizing current

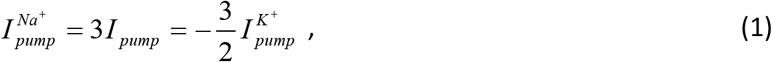

in response to the intracellular [Na^+^] (as per **Figure 9**). In CN-CD only there is a K^+^/Cl^-^ co-transporter for ion flow, *J*_*KCl*_.

Since animal cells cannot sustain osmotic pressures, intra/extracellular osmolyte concentration inequalities produce a H_2_O flow till osmotic balance is restored; this results in Vol_cell_ changes at rates set by the smaller of the two components giving rise, at any time, to osmotic loading, i.e. to net [Na^+^ + Cl^-^] entry.

### Choice of leak permeabilities

Whereas CN-CD is precisely the Dijkstra et al (2016) model, the SMF version (SM-CD) has a leak permeability ratio 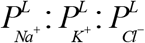 broadly consistent with Fraser and Huang (2004); for more detail see **Setting V**_**m**_ below.

### GHK driving forces

Currents through open channels (permeability pathways), ion-specific or not, are modeled with the GHK formulation (Hille, 2001). For ion X = Na^+^, K^+^, or Cl^-^, the GHK current is given by:

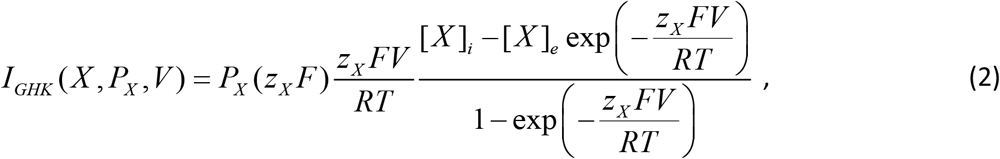

where *P*_*X*_ is the permeability, *z*_*X*_ the valence, and [X]_i_ and [*X*]_e_ the intra- and extra-cellular concentrations of *X* respectively.

### Leak currents

“Leak” permeability mechanisms use the GHK formulation:

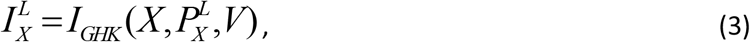

where *X* is either *Na*^*+*^, *K*^*+*^ or *Cl*^*-*^. Leak permeability 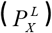 values were adjusted for model variants used here (**Table 2**) in the context of appropriate setting of *V*_*rest*_.

### Cation channel currents

For non-selective cation channels, we use a P_K_:P_Na_ ratio of 1:1.11 and the formulation:

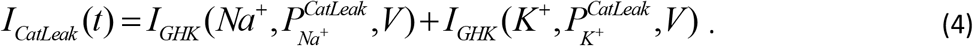

In the present models, cation channel leaks do not contribute to healthy steady-states – they are either transient stimulatory currents through SMF-endplate type AChR channels, or pathological leaks (hence “leaky” cation channels).

#### Transient voltage-gated Na^+^ current

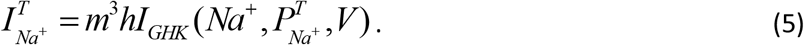

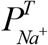is the maximal membrane permeability to Na^+^ through a V-gated channel (operating in a Hodgkin-Huxley (H-H) fashion). *m* is the H-H Na^+^ channel activation/deactivation gating variable and *h* is the H-H Na^+^ channel inactivation/recovery gating variable. The current’s driving force also follows the GHK form of **Equation 1**.

### Delayed rectifier K^+^ current

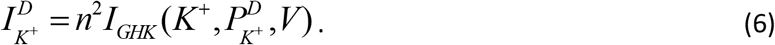

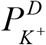 is the maximal membrane permeability of K^+^ through a V-gated channel (operating in a H-H fashion). *n* is the delayed rectifier K^+^ channel activation/deactivation gate variable.

### Voltage-dependent gating

The non-dimensional gating parameters *m, h, n* evolve in time according to:

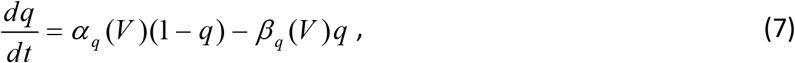

where *q* ϵ {*m, h, n*} are as defined above, and the voltage dependent α_q_(*V*) and β_q_(*V*) refer, for *m and n* to gate activation and deactivation and, for *h* to inactivation and recovery from inactivation. **Table 4** gives the voltage dependences for the relevant rate constants.

### Voltage-dependent Cl^-^ current

The SLC26A11 ion exchanger based voltage-dependent Cl^-^ conductance (Rungta et al, 2015) is as described by Dijkstra et al (2016):

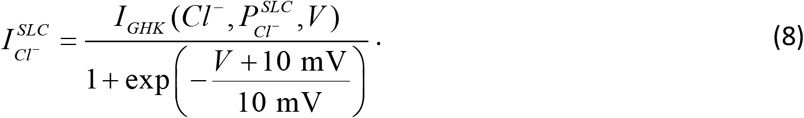

Except in CN-CD and MN-CD, this pathological cortical neuron specific conductance is set at zero.

### K^+^/Cl^-^ cotransporter

This cotransporter (strength *U*_*KCl*_) is present only in CN-CD (it is not evident in skeletal muscle; Pedersen et al 2016) where (as per Dijkstra et al 2016) it is given by

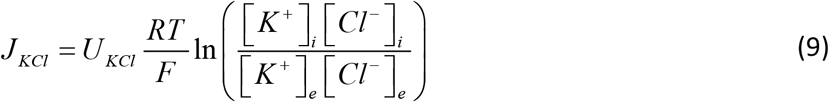

(except that there is an error in the log term of Eq. (6) in Dijkstra et al 2016).

### 3Na^+^/2K^+^ ATPase pump current

The electrogenic 3Na^+^(out)/2K^+^(in) ATPase modeled here is that of Hamada et al (2003):

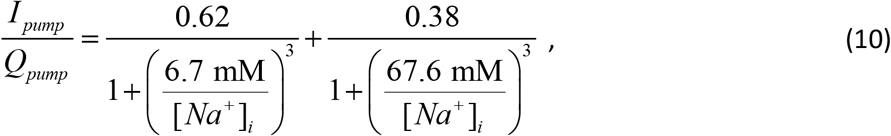

where

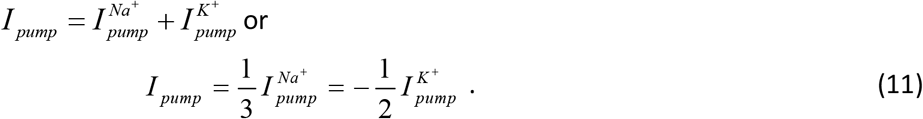

Thus *I_pump_* signifies a net hyperpolarizing Na^+^ outflow. The Hamada et al (2003) formulation (**Figure 9**) is appropriate for models that assume an invariant extracellular medium. It depicts the pump’s two intracellular Na^+^-binding sites (with ∼10-fold different affinities) but has no term for the extracellular K^+^*-* binding site. Here, pump-strength (*Q*_*pump*_) is varied in many computations (e.g., it is set to zero to depict anoxia, diminished towards zero to depict ischemia and multiplied for up-regulation).

### ATP-Consumption

ATP-consumption by the Na^+^/K^+^ ATPase pump in the all the CD models is given as

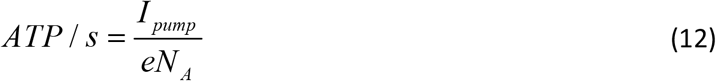

where *e* is the electronic charge (1.6 x 10^−19^Coul) and *N*_*A*_ is the Avogadro number. In other words, *I*_*pump*_ of 1 pA is equivalent to an ATP-consumption of 10.38 amol/s.

### Cell Volume

Cellular swelling is driven by an influx of water at rates that depend on the transmembrane osmotic gradient. The rate of change of cell volume, Vol_cell_, due to water influx is given by:

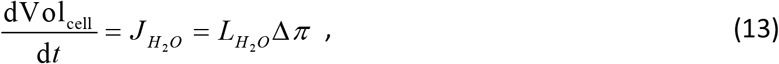

where Δ*π RT*([*S*]_*i*_ − [*S*]_*e*_), and [*S*]*i* and [*S*]*e* denote total concentrations of intra- and extracellular solutes and 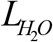 is the effective membrane water permeability. This equilibration is typically assumed to be nearly instantaneous relative to the ion flows (Fraser and Huang, 2004; Dijkstra et al 2016; Kay, 2017; though see also Dmitriev et al 2019). Accordingly, in modeling here, osmotic ion fluxes, not the H_2_O fluxes what limit the rate of osmotic swelling or shrinkage.

### Nernst potentials

The equilibrium potential (Nernst potential) for each ion

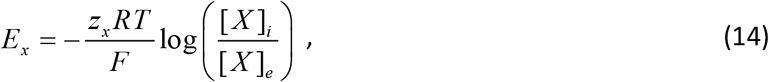

where *X* is either Na^+^, K^+^, or Cl^-^ ions and *zX* is the valence of each ion.

### Charge difference and membrane potential

Because CD models keep track of the absolute number of ions flowing across the cell membrane (of capacitance *C*_m_) no differential equation is needed for *V*_m_. Instead, an accounting equation is used, made simple because the extracellular space is kept neutral:

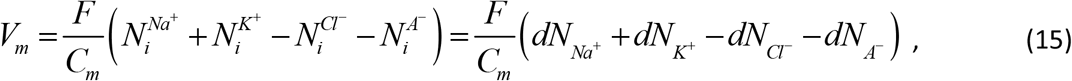

where 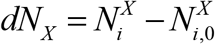 is the difference in the number of ions of species *x*, between its present value 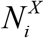and a reference value 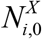. The reference values are those yielding *V*_*m*_ = 0 V, which is equivalent to a neutral intracellular space (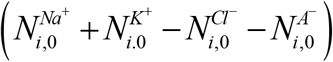). Because N_A-_ is constant, note that 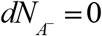 in **Equation 15**. Number differences, *dNx*, are easier to work with than total numbers of ions *N*_*x*_ (or than concentrations). For typical cell C_m_ values, voltages in the mV range correspond to net intracellular charge differences (measured as a number of singly charged ions) of the order of attomoles (10^−18^ moles). For instance, when *V*_*m*_ varies in the range [-100mV, 100mV] this corresponds to a change in net singly charged ions of [-20.7, +20.7] amol. With our *Vol*_cell_ = 2000 µm^3^, this yields tiny concentration changes [-0.01, 0.01] mM. For solutions whose concentrations are in the 1-100 mM range, there could be 4-5 orders of magnitude difference between steady-state values and the changes (in the 0.0001 to 0.01 mM range). Computations based on concentrations, therefore, tend to be unstable. For this reason, calculations in this study are based on changes in ion numbers (in amol units).

### Number of intracellular ions

CD models account for the change in intracellular ions at any given moment (Fraser et al 2004; Dijkstra et al 2016), via the following simple relationships of the respective currents:

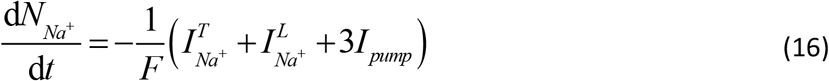

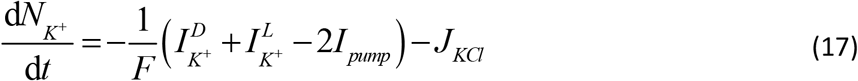

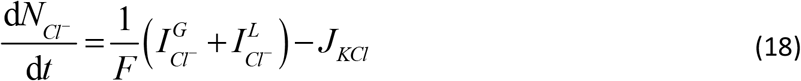

### Setting V_rest_ in SM-CD and other CD models

Experimentally, excitable cells’ *V*_*rest*_ (i.e., steady-state *V*_*m*_) values are generally more accessible than cytoplasmic ion concentrations or cell volume, making it useful to anchor a model with a consensus *V*_*rest*_ value. For SM-CD, we chose V_rest_= −86, with parameter determinants established in an iterative process as follows: first, a number is chosen for impermeant anions 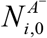, consistent with the system’s total cation concentration (as per the extracellular solution). We started with the CN-CD value, knowing that fine-tuning would be needed to meet our (self-imposed) requirement that SM-CD have the same resting Vol_cell_ as CN-CD. Thus, note in **Table 2** the slightly different 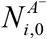 for CN-CD and SM-CD (likewise, LA-CD and SM-CD, to give identical resting Vol_cell_).

At rest, Na^+^ and K^+^ leak currents (for a system with a given pump-strength), must precisely balance. In other words, *V*_*m*_ converges (along with ion concentrations and Vol_cell_) on steady-state (=*V*_rest_) when

*passive Na*^+^ influx + *active* Na^+^outpump = zero

AND

*passive K*^+^ efflux + *active* K^+^inpump = zero.

For the 3Na^+^/2K^+^ ATPase pump current, this requirement is met when:

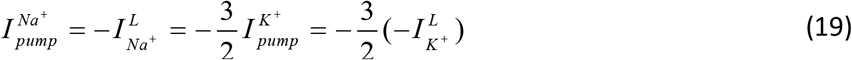

For P-L/D systems with a given pump-strength, *V*_*rest*_ varies monotonically with P_Na_:P_K_ ratio; low ratios yield hyperpolarized *V*_*rest*_ values, high ratios, depolarized ones (**Figure 10**). *V*_*rest*_=-86 mV for SM-CD requires P_Na_:P_K_ = 0.03 : 1 (likewise for WD-CD). Absolute values for P_Na_ and P_K_, and for P_Cl_ (**Table 2**) were guided by the P_Na_:P_K_:P_Cl_ ratio (0.02: 1: 3) reported for amphibian skeletal muscle (Fraser and Huang 2004). For absolute P values, a straightforward constraint was the choice to give SM-CM the same area (C_m_) as CN-CD.

As per **Equation 16**, a pump stoichiometry other than 3Na^+^out/2K^+^in would, all else being equal, alter V_rest_. Pump stoichiometry is invariant here, but see Dmitriev et al (2019).

### Excitability and safety factor for SM-CD

For SM-CD to be hyperpolarized and appropriately excitable (i.e. “relatively inexcitable”), its input impedance had to be a) notably less than for CN-CD and b) predominantly P_Cl_-based (Pedersen et al 2016). With resting-P values set, the need to trigger spikes near −60 mV (Fu et al 2011) with a reasonable-sized safety factor (Ruff 2011) had to be met. To achieve safety factor ∼1.5, Nav and Kv “densities” in SM-CD were set at 3X the CN-CD level (a larger P_Cl_ would have required even greater V-gated channel densities). Thus, absolute P_Cl_, P_Na_ and P_K_ values of SM-CD are biologically appropriate, but leave room for physiological modulation (to, say, alter V_rest_ via ΔP_Na_ or ΔP_K_, or to modulate excitability by ΔP_Cl_).

**Table 2** shows that m^3^h is vanishingly small at V_rest_ in SM-CD; for CN-CD it adds an extremely small Nav channel contribution to the operational value of P_Na_ that negligibly affects V_rest_.

### Cytoplasmic Donnan effectors

Once V_rest_ is set via P_Na_ and P_K_, the intracellular anion concentrations are determined uniquely for the resting state. If Cl^-^ transport is purely passive (i.e., no involvement of secondary transport) as in SM-CD, then E_Cl_ = V_rest_ and therefore:

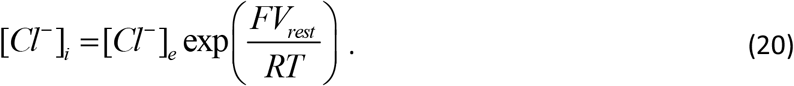

[A^-^]_i_ then follows from the voltage and osmotic balance requirements. The voltage requirement yields:

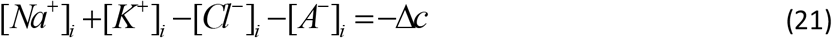

where Δc is the tiny excess concentration of anions associated with V_rest_ and equal to [intracellular anions]-[intracellular cations]:

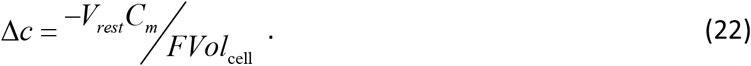

Δc = 0.00886 mM for *V*_*rest*_ = −86 mV. And the osmotic equilibrium condition is:

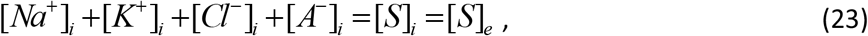

where [S]_I_ and [S]_e_ are defined below **Equation 13. Equations 21** and **23** yield:

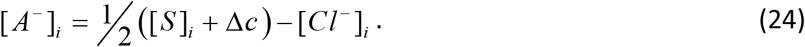

A P-L/D model-cell’s design for steady-state imposes its [A^-^]_i_. Cell volume adjusts to reflect that constraint 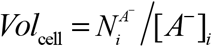. As mentioned above, SM-CD’s 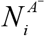 was set to give resting Vol_cell_ =2000 μm^3^, i.e. 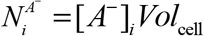. Here, the quantity of impermeant anions 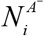 is invariant, but for many *in vivo* circumstances, it would vary, and variations in Vol_cell_ would be expected in such cases.

### Instantaneous perturbations

Experimental solution changes (e.g., as in brain slice experiments) typically require finite “wash-in/wash-out” times. Dijkstra et al (2016) mimicked such solution changes (affecting pump rates and channel gating etc) but here, doing so would have unnecessarily obscured mechanistic underpinnings of responses. Thus, pump-off (anoxia) and pump-on (restoration of pump-strength) changes and channel gating changes (Nav and cation channels open probabilities) are instantaneous.

### Maximum cell volume before lysis

Both CN-CD and SM-CD have C_m_=20pF and steady-state Vol_cell_=2000 µm^3^. If 0.01F/m^2^ (= 0.01 pF/µm^2^) is the specific capacitance of the bilayer, membrane area is 20/0.01 = 2000 µm^2^. With 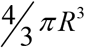 the volume of a spherical cell and 4π*R*^*2*^ its surface area, maximum Vol_cell_ as the cell swells (to spherical) would be =4/3π(2000/4π)^3/2^ = 8410.4 µm^3^. Given a 4% bilayer elasticity strain limit (yielding membrane area = 2080 µm^2^), rupture would occur at 8920 µm^3^. Thus, in bifurcation plots, the notional Donnan Equilibrium (DE) values indicated for reference, are unachievable by these models. Note too that present models depict neither surface area regulation nor membrane tension homeostasis (see Morris 2018).

### Excitatory post-synaptic current (EPSC) via AChR channels

SM-CD action potentials are initiated by EPSCs, i.e. macroscopic end-plate currents through (acetylcholine receptors) AChRs, which are non-selective cation channels that pass Na^+^ and K^+^ as per the GHK formalism. The EPSC time course mimics the *g* (*t*) reported in Wang et al (2004)(see **Figure 1A top**). As reported in Hille (2001), 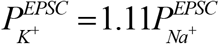. The function *g*(*t*) has a maximum of 1 and 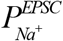 has a 1.5-fold safety factor (i.e., an amplitude adjusted to 1.5X the threshold required to elicit an AP in SM-CD). The end-plate current is:

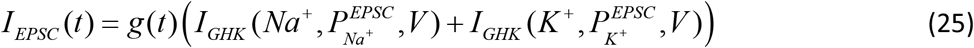

### Computational methods

Calculations involved solving sets of first order differential equations. These were done using Python with the ordinary differential equation solver *odeint*.

## ACKNOWLEDGEMENTS

We acknowledge financial support from Natural Sciences and Engineering Council (Canada) (BJ and JJW) and support from the Ottawa Health Research Institute (CEM).

## Supplemental Figures

**FIGURE Sup 1.**
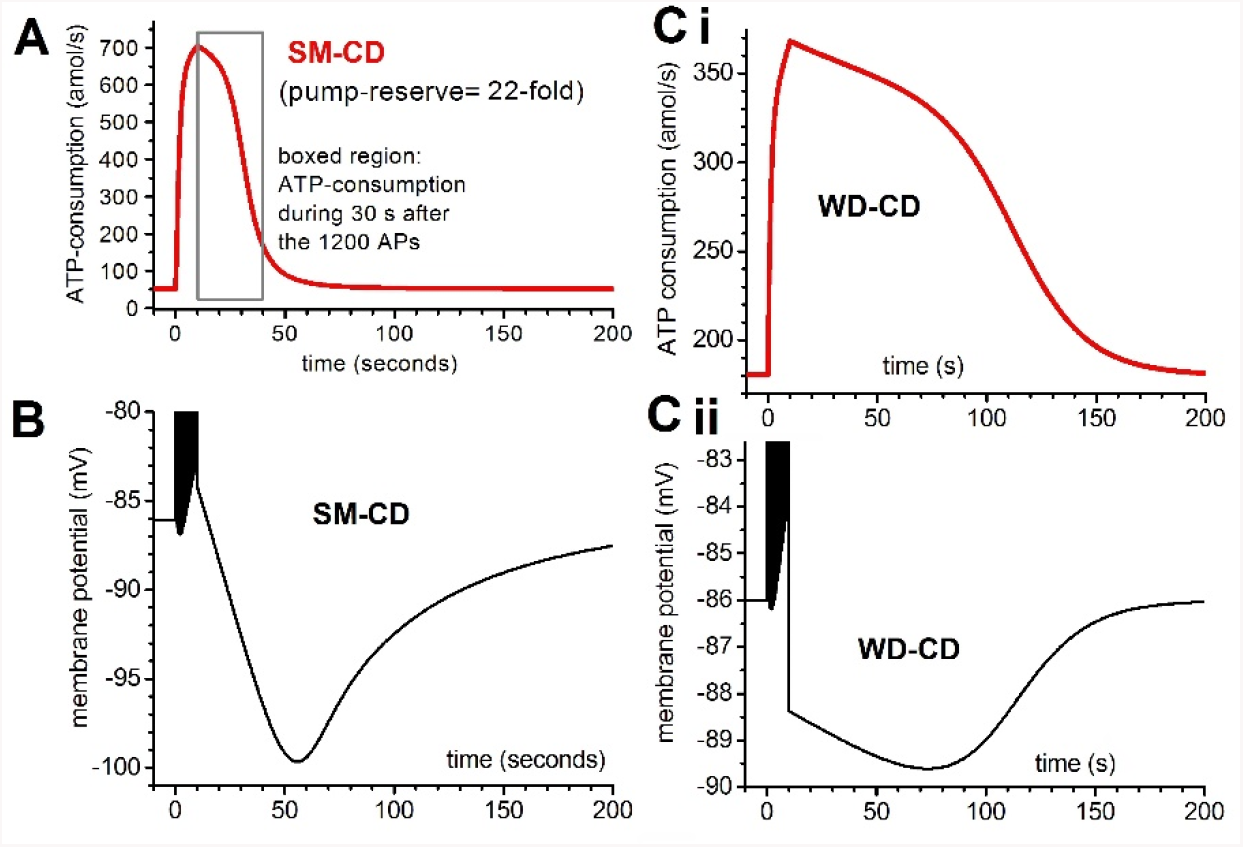
Stress-test. **A**. By ∼30 seconds after APs stop in SM-CD, ATP-consumption (Na^+^-extrusion) it almost back to steady-state when pump-strength=200% (maximum ATP-consumption = 2×566 = 1,132 amol/s or I_pumpmax_ 2×54.5 = 109 pA) i.e., when pump-reserve= 22.2-fold. **B**. V_m_(t) for SM-CD (pump-strength 100%) from **Figure 2A** but expanded to emphasize the effect of hyperpolarizing I_pump_ during ion homeostatic recovery, post-1200 APs. **Ci**. For the same 1200 AP stress-test performed by SM-CD (**Figure 2A**), ATP-consumption by WD-CD. **Cii**. V_m_(t) for WD-CD expanded over the voltage range relevant to ion homeostatic recovery. At 10 seconds (immediately post-1200 APs) V_m_ goes to −88.5 mV (then, to −89.8 mV before returning to steady-state). Also at t=10 seconds, ATP-consumption peaks at 370 amol/s while (not shown), [Na^+^]_i_=32 mM, [Cl-]i=5.75 mM, and Vol_cell_ =2004.5 µm^3^.

**FIGURE Sup 2.**
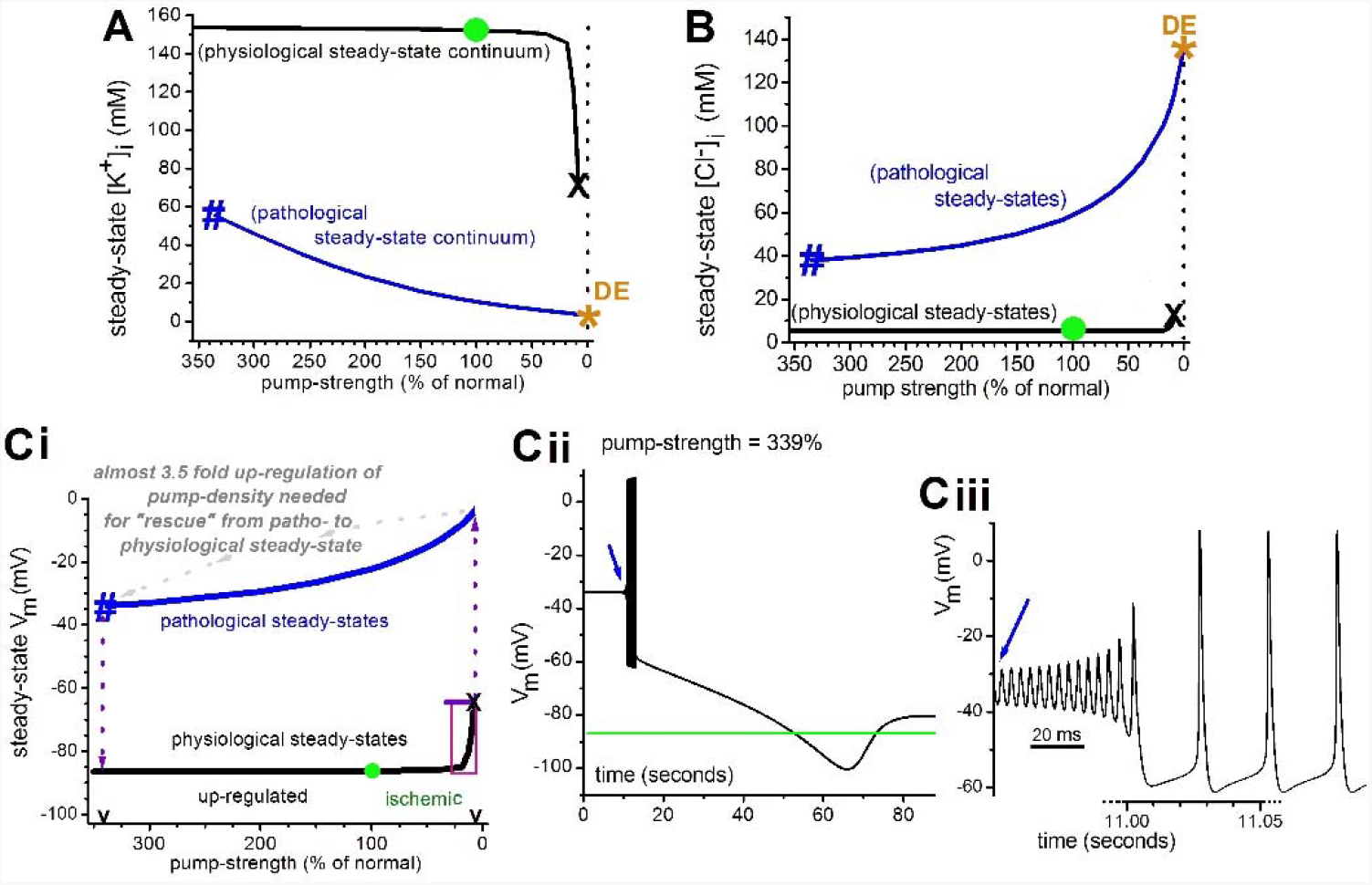
Bifurcation analysis. **A,B**, from the same SM-CD bifurcation analysis as in **Figure 7A,B,C** plots for steady-state [K^+^]_i_ and [Cl^-^]_i.._ **Ci**, the SM-CD V_m_ bifurcation plot showing the location of **#**, the unstable threshold on the pathophysiological steady-state continuum. The equivalent bifurcation point in CN-CD is called a Hopf bifurcation by Dijkstra et al (2016). **ii,iii** At t=0, SM-CD (on the pathological continuum) pump-strength is stepped from 338% to 339%. The resulting trajectory is plotted (**ii**), with a portion expanded (**iii**).

